# Light chain 2 is a Tctex-type related axonemal dynein light chain that regulates directional ciliary motility in *Trypanosoma brucei*

**DOI:** 10.1101/2021.09.29.462335

**Authors:** Subash Godar, James Oristian, Valerie Hinsch, Katherine Wentworth, Ethan Lopez, Parastoo Amlashi, Gerald Enverso, Samantha Markley, Joshua Alper

## Abstract

Flagellar motility is essential for the cell morphology, viability, and virulence of pathogenic kinetoplastids, including trypanosomes. *Trypanosoma brucei* flagella exhibit a bending wave that propagates from the flagellum’s tip to its base, rather than base-to-tip as in other eukaryotes. Thousands of dynein motor proteins coordinate their activity to drive ciliary bending wave propagation. Dynein-associated light and intermediate chains regulate the biophysical mechanisms of axonemal dynein. Tctex-type outer arm dynein light chain 2 (LC2) regulates flagellar bending wave propagation direction, amplitude, and frequency in *Chlamydomonas reinhardtii*. However, the role of Tctex-type light chains in regulating *T. brucei* motility is unknown. Here, we used a combination of bioinformatics, in-situ molecular tagging, and immunofluorescence microscopy to identify a Tctex-type light chain in the procyclic form of *T. brucei* (TbLC2). We knocked down TbLC2 expression using RNAi, rescued the knockdown with eGFP-tagged TbLC2, and quantified TbLC2’s effects on trypanosome cell biology and biophysics. We found that TbLC2 knockdown resulted in kinetoplast mislocalization and the formation of multiple cell clusters in cell culture. We also found that TbLC2 knockdown reduced the directional persistence of trypanosome cell swimming, induced an asymmetric ciliary bending waveform, modulated the bias between the base-to-tip and tip-to-base beating modes, and increased the beating frequency. Together, our findings are consistent with a model of TbLC2 as a down-regulator of axonemal dynein activity that stabilizes the forward tip-to-base beating ciliary waveform characteristic of trypanosome cells. Our work sheds light on axonemal dynein regulation mechanisms that contribute to pathogenic kinetoplastids’ unique tip-to-base ciliary beating nature and how those mechanisms underlie dynein-driven ciliary motility more generally.

**Author Summary:** Kinetoplastea is a class of ciliated protists that include parasitic trypanosomes, which cause severe disease in people and livestock in tropical regions across the globe. All trypanosomes, including *Trypanosoma brucei*, require a cilium to provide propulsive force for directional swimming motility, host immune evasion, and various aspects of their cell cycle. Thus, a functional cilium is essential for the virulence of the parasite.

Trypanosome cilia exhibit a unique tip-to-base beating mechanism, different from the base-to-tip beating of most other eukaryotic cilia. Multiple ciliary proteins are involved in the complex biophysical and biochemical mechanisms that underly the trypanosome ciliary beating. These include dynein motor proteins that power the beat, dynein-related light chains that regulate the beat, and many other proteins in the nexin-dynein regulatory complex, in the radial spokes, and associated with the central pair of microtubules, for example.

Here, we identify a Tctex-type dynein light chain in *T. brucei* that we named TbLC2 because it has sequence homology, structural similarity, and ciliary localization like LC2 homologs in other organisms. We demonstrate that TbLC2 has critical dynein regulatory functions, with implications on the unique aspects of trypanosome ciliary beating and cellular swimming motility. Our study represents an additional step toward understanding the functions of the trypanosome ciliary proteome, which could provide novel therapeutic targets against the unique aspects of trypanosome ciliary motility.

## Introduction

Motile cilia and flagella (terms used interchangeably, here we favor cilia) are multifunctional and dynamic eukaryotic cell organelles that drive cell motility, generate fluid flows across surfaces, sense mechanical and chemical signals, and serve as cellular adhesion and secretion sites [1]. In *Trypanosoma brucei*, a pathogenic Kinetoplastea class member that causes African Trypanosomiasis in humans and nagana in livestock, has a single motile cilium that runs laterally along the length of the cell. Ciliary motility is essential throughout the life cycle of the parasite [2]. Trypanosomes use their cilium to generate propulsive force for directional swimming, proper morphogenesis, completion of cytokinesis, and initiation of kinetoplast (a network of interlocked circular DNA, kDNA, inside a large, single mitochondrion associated with the basal body [3]) segregation during cell division [4]. Additionally, ciliary motility plays a significant role in tsetse fly host midgut epithelial layer attachment, migration from the midgut to the salivary gland, and evasion of the immune system in mammalian hosts [5]. Therefore, proteins enabling and regulating ciliary motility are vital for parasite viability and pathogenicity [6], and, through the combination of proteomic [7] and molecular mechanistic studies, ciliary proteins essential for motility could provide an excellent prospect for novel therapeutic drug targets [2].

Trypanosome cilia generate motile force using the axoneme, a highly-conserved structure made of a “9+2” arrangement of microtubule doublets, dynein motor proteins, and approximately 750 other ciliary proteins [7]. Inner (IAD) and outer arm dyneins (OAD) [8] are minus-end-directed motor protein complexes [8] comprising multiple heavy chains (HCs), intermediate chains (ICs), and light chains (LCs) that power ciliary motility. Dynein heavy chain tail domains permanently attach dynein to one microtubule doublet while its microtubule-binding domain (MTBD) binds to the adjacent doublet [9] and slides it distally in an ATP-dependent manner [10, 11]. The nexin-dynein regulatory complex (N-DRC) and basal constraints convert doublet sliding to bending [12]. Coordination mechanisms that activate [13] and deactivate [14] dynein along and across the axoneme ultimately orchestrate a bending wave that is necessary for motile functionality [15]. In *Trypanosoma brucei* and other Trypanosomatidae cells, this bending wave propagates from the tip to the base of the cilium [2], which is different from the canonical base-to-tip bending wave propagation direction found in nearly all other eukaryotic cilia and flagella.

Wild-type *T. brucei* cells exhibit highly directionally persistent swimming motility due to their tip-to-base propagating ciliary beat waveform [16]. However, a tumbling maneuver transiently interrupts their directionally persistent swimming motility mode [16]. An abrupt switch to a base-to-tip propagating ciliary beat waveform initiates tumbling [17] and reorients the swimming direction. While less than 25% of the wild-type population demonstrate the tumbling mode [16, 18], trypanosome cells with mutations to the N-DRC tumble rather continuously [4, 19], suggesting that dynein motor regulation underlies tip-to-base ciliary beating in the directionally persistent swimming motility mode of trypanosome cells and the switch to the base-to-tip driven tumbling motility mode.

There is significant additional evidence from multiple species supporting the model of axonemal dynein motors requiring significant regulation [10] to achieve the precise and regular tip-to-base ciliary beat waveforms necessary for persistent directional motility [20]. Inner and outer arm axonemal dynein regulation acts through multiple direct and indirect interactions, including through their ICs and LCs, which respond to various stimuli, including Ca^2+^, phosphorylation, and redox poise [21–23], and in conjunction with the N-DRC, radial spokes, and the central pair complex [11, 12]. For example, LC4, a calmodulin-like outer arm dynein light chain also known as DLE1 [13], induces a Ca^2+^ dependent conformational change in the N-terminal tail domain of the γ HC (DHC15 [13]) that activates the motor in *Chlamydomonas reinhardtii* cells (a single-celled alga that is commonly used as a model organism for ciliary motility) [22]. The loss of LC4 from *Leishmania mexicana*, another pathogenic Kinetoplastea, results in a higher ciliary beat frequency and an increased swimming speed [24]. LC1, a leucine-rich repeat dynein light chain, directly interacts with the MTBD of γ HC in *Chlamydomonas* cells [14]. The loss of LC1 from *C. reinhardtii* and *T. brucei* changes the ciliary beat frequency and beating mode, which reverses the direction of motility [20, 25].

LC2 is a member of the Tctex-type family, and it is essential for OAD assembly in *Chlamydomonas* [26]. However, *Chlamydomonas* cells with an N-terminal truncation on LC2 have partially functional cilia with the OADs still intact [9], demonstrating that mutations which affect the dynein regulatory function of LC2 can cause ciliary motility defects separate from the structural defects caused by the complete loss of the protein. LC2 has no direct interactions with the dynein HCs during any stage of their mechano- chemical cycle [20], yet it exhibits OAD regulatory function [9], which suggests possible LC-LC or LC-IC interactions may be necessary for LC2 to carry out its regulatory function. Additionally, inactivation of Tcte3-3, an LC2 homolog in mice, causes motility defects in sperm cells characterized by reduced motility persistence [21]. Inactivation of Tcte3-3 also increased the ciliary waveform’s beat frequency and decreased its amplitude in spermatozoa, rendering them unable to migrate into the female oviduct [21]. Similar to the *Chlamydomonas* mutant with LC2 N-terminus truncation [9], murine cilia missing Tcte3-3 showed no structural defects within the axoneme, including intact OADs [21], but some spermatozoa had two cilia, and others showed a bent or coiled phenotype [21]. Moreover, LC2 is phosphorylated upon the activation of rainbow trout and chum salmon sperm motility [22], and reduced phosphorylation of LC2 correlates with less progressive swimming of the spermatozoa [22]. In total, these results suggest that LC2 is an essential regulator of axonemal dynein function and show its loss or mutation leads to ciliary motility defects. However, a quantitative functional analysis of the roles played by LC2 in regulating the ciliary motility, cell morphology, and ultimately in the swimming behavior of Kinetoplastea, including disease-causing parasites like *T. brucei*, is lacking.

In this study, we identified a trypanosome homolog (Tb927.9.12820, hereafter called TbLC2) of the *Chlamydomonas* Tctex1/Tctex2 family outer arm dynein light chain 2. We knocked down the expression of endogenous TbLC2 in *T. brucei* cells using RNA interference and rescued the knockdowns by overexpressing a recombinant fluorescent marker (eGFP) tagged TbLC2 (TbLC2::eGFP). We found that TbLC2 knockdowns exhibited cell division and cell culture growth rate defects. We also found that TbLC2 knockdowns had reduced directional swimming persistence characterized a higher beat frequency, ciliary bending waveform defects, and a shift in the waveform propagation direction bias from the tip-to-base to the base-to-tip beating mode. Because knocking down TbLC2 did not cause significant structural phenotypes in the trypanosome cilia but did cause the functional phenotypes similar to, however distinct from, other species, our data suggest that TbLC2 regulates axonemal dynein and the tip-to-base beating waveforms characteristic of *Trypanosoma brucei* ciliary motility.

## Results

### TbLC2 shows high sequence and structural conservation with outer arm dynein Tctex-type light chains

We searched for Tctex-type dynein light chains in the *Trypanosoma brucei* genome (TriTrypDB [27]) using the *C. reinhardtii* outer arm axonemal dynein light chain 2 (Cre12.g527750 in *C. reinhardtii* v5.6 [23, 28], which we refer to as CrLC2) as the query sequence in protein BLAST [24]. We identified Tb927.9.12820 and Tb927.11.7740 as two putative Tctex-type dynein light chain homologs (Materials and Methods). We cross-referenced these two hits for ciliary localization in the TrypTag database [29], and we found that only Tb927.9.12820, previously annotated as a putative dynein light chain [27], strongly localized to the cilium. Therefore, hereafter we refer to Tb927.9.12820 as *Trypanosoma brucei* dynein light chain 2 (TbLC2).

We aligned TbLC2 with LC2 proteins from multiple species (protozoans to humans, S1 Fig). We found 29% sequence identity over a 124-amino acid region of overlap between *C. reinhardtii* and *T. brucei* (S1 Fig). We also identified significant extensions in the N-terminal domain of mouse (*Mus musculus*) and rainbow trout (*Oncorhynchus mykiss*) LC2, as well as gaps in the N-terminal regions of *T. brucei* and *Trypanosoma cruzi* LC2 (S1 Fig).

We generated sequence-based structural homology and template-free models of TbLC2 using SWISS-MODEL (S1 File) [30] and AlphaFold (S2 File) [31] (Materials and Methods), respectively. We compared the cryo-EM structure of CrLC2 in the outer arm dynein core subcomplex from *C. reinhardtii* (PDB ID: 7KZN [20]) to these models (Fig 1A). We found a high degree of structural alignment between CrLC2 and our homology and template-free models of TbLC2 (Fig 1A, RMSD = 0.1 Å and 1.7 Å, respectively). We also found a high degree of local surface charge density conservation between CrLC2 and the models of TbLC2 (Fig 1B-D, lower panels), particularly in the suggested binding interface between CrLC2 and CrLC9 (S2 Fig) [20]. Of the 35 residues in CrLC2 constituting the suggested binding interface (S2 Fig), 12 residues were identical and 20 were hydrophobicity and polarity preserving substitutions (S2 Fig). In particular, R100 and R104 in CrLC2, which constitute 3/9 of the polar contacts with the adjacent CrLC9, were conserved (identical, R88, and charge conserving, K92, respectively) in TbLC2 (Fig 1A and S2 Fig).

**Fig 1.**
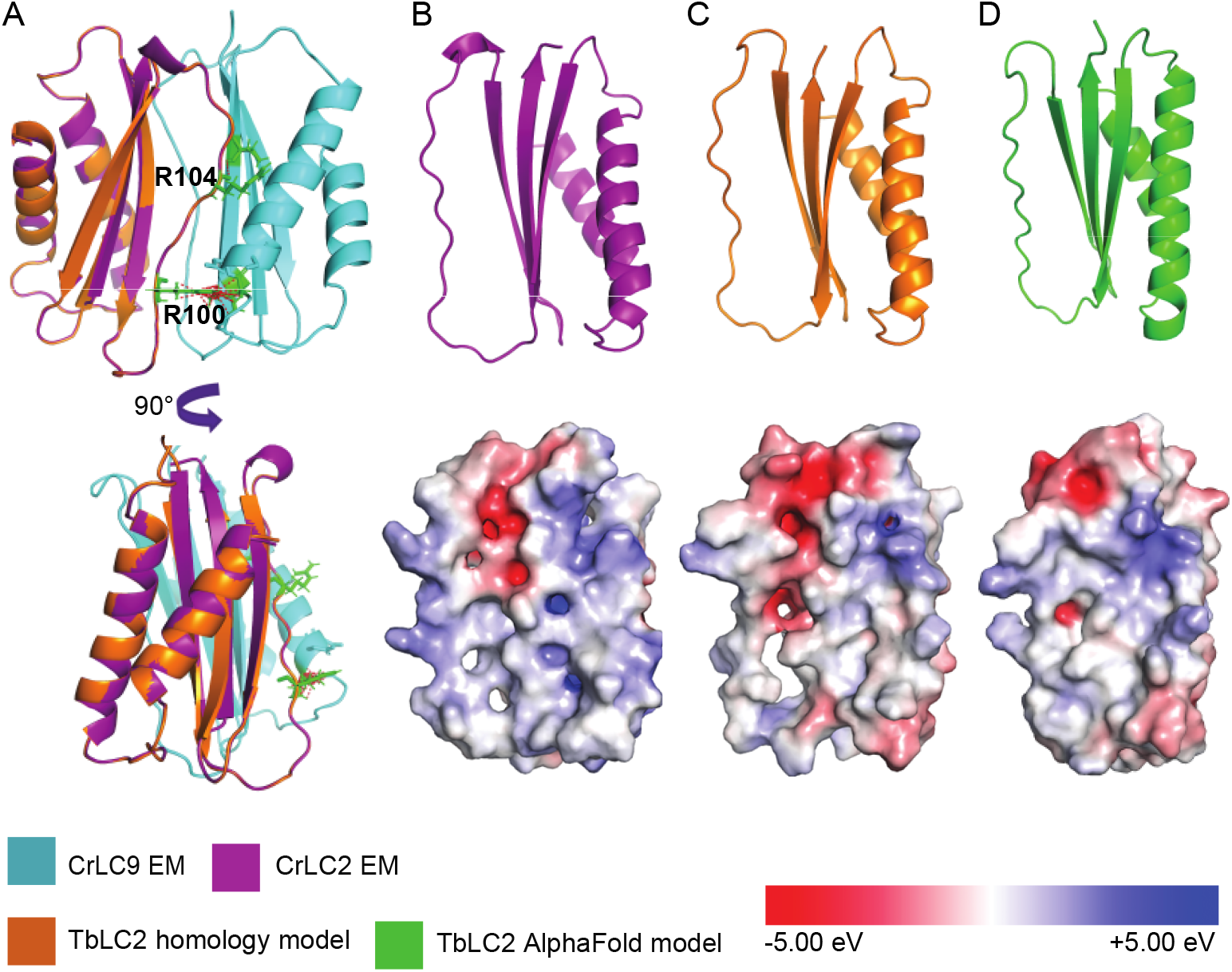
TbLC2 shows a high degree of structural and electrical potential conservation with the *C. reinhardtii* LC2 protein. **A.** Alignment of the sequence-based structural homology model of TbLC2 (*orange*) with LC2 and LC9 (denoted as CrLC2, *purple,* and CrLC9, *cyan*, respectively) from the *C. reinhardtii* outer dynein arm core subcomplex structure (PDB ID: 7KZN[20]). R100 and R104 (*green*) in CrLC2, which are highly conserved polar residues, have extensive contacts (*red dotted line*) with CrLC9. The structures (*top*) of CrLC2 in **B.,** the structural homology model of TbLC2 in **C.,** and the template-free model of TbLC2 in **D.**, are shown with surface charge density mapped to the electrostatic potential distribution (*bottom*).

### TbLC2 localizes along the axoneme

To examine the cellular localization of TbLC2 in trypanosome cells, we overexpressed recombinant, eGFP-tagged LC2 (TbLC2::eGFP) in wild-type procyclic-form trypanosome cells (hereafter called WT/LC2 OE cells, Table B in S1 Text for *T. brucei* strains generated and used in this study, Material and Methods). We verified the expression of the TbLC2::eGFP in WT/LC2 OE cells using flow cytometry (Material and Methods) and found that WT/LC2 OE cells had a significantly higher eGFP fluorescence intensity than the background fluorescence observed in the parental wild-type cells (S3 Fig). We also imaged the WT/LC2 OE cells (Material and Methods) and found high levels of eGFP fluorescence throughout the cells (Fig 2A). To confirm ciliary localization of TbLC2, we immunostained the cells (Materials and Methods) using a primary antibody against paraflagellar rod 2 protein (PFR2 [6]). We observed TbLC2::eGFP (Fig 2A, *green*) and PFR2 (Fig 2A, *red*) signals along most of the cilium’s length, apart from the tip where the PFR2 signal ran shorter than the eGFP signal (Fig 2A) suggesting the ciliary localization of TbLC2. However, the high cytoplasmic fluorescence signal prevented us from conclusively confirming the localization of TbLC2 to the cilium because it was attached to the cell body (Fig 2A).

**Fig 2.**
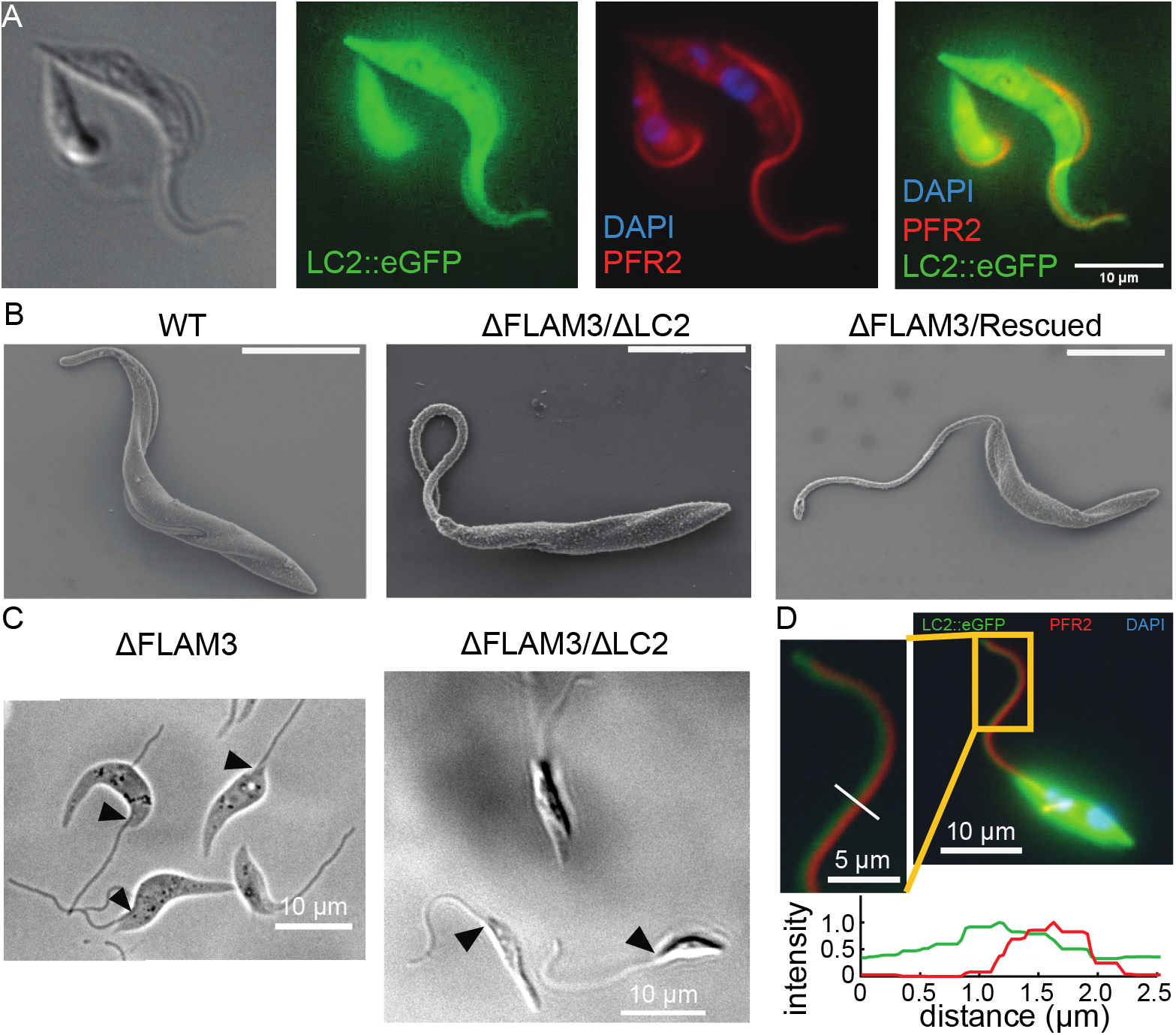
Overexpressed TbLC2 localized to the axoneme as well as the cytoplasm. **A.** Doxycycline induced TbLC2::eGFP overexpressing wild-type cells (WT/LC2 OE cell) visualized in DIC (*left*) and under widefield fluorescence microscopy (*right*). The TbLC2::eGFP (*green*) localized to the cytoplasm and the cilium. The immunostained paraflagellar rod 2 (PFR2, *red*), DAPI stained DNA (DAPI, *blue*), and TbLC2::eGFP overlay (*right*), suggest that the TbLC2 has ciliary and cytoplasmic localization. **B.** SEM images showing cilium and cell body morphology of wild-type (WT, *left*), ΔFLAM3/ΔLC2 (*middle*), and ΔFLAM3/Rescue (*right*) cells. The scale bar is 5 µm in each panel. **C.** ΔFLAM3 (*left*) and ΔFLAM3/ΔLC2 (*right*) cells visualized with DIC microscopy showing short flagellar attachment zones (*arrows*) and nearly complete ciliary detachment from the cell body. **D.** Doxycycline TbLC2::eGFP expressing ΔFLAM3/Rescue cells stained with DAPI (*blue*) and the anti-PFR2 antibody (*red*) visualized under a widefield fluorescence microscope. The inset shows that TbLC2::eGFP and the paraflagellar rod are separately localized, and the line scan plots shows the normalized fluorescence intensity of TbLC2::eGFP (*green*) and PFR2 (*red*) along the line (*white*) shown in the inset.

We used RNA interference to knock down FLAM3, a flagellar attachment zone protein, to detach the cilium from the trypanosome’s cell body (ΔFLAM3 cells, Materials and Methods [32]). Additionally, we knocked down endogenous TbLC2 expression using RNAi in wild-type (hereafter called ΔLC2 cells, Materials and Methods) and ΔFLAM3 cells (hereafter called ΔFLAM3/ΔLC2 cells, Materials and Methods) to improve the efficiency of expressed TbLC2::eGFP assembly into the cilium (hereafter called ΔFLAM3/Rescued cells, Materials and Methods). We found that the FLAM3 knockdown in ΔFLAM3/ΔLC2 cells resulted in the cilia being mostly separated from the trypanosome cell bodies (Fig 2B and Fig 2C), as previously reported [32]. With the cilium detached from the cell body, we detected eGFP fluorescence along the cilia of ΔFLAM3/Rescued cells (Fig 2D). To further isolate the sub-ciliary localization of TbLC2::eGFP and PFR2, we immunostained ΔFLAM3/Rescue cells using a primary antibody against PFR2 (Materials and Methods). We observed a fixed, finite separation (300 - 400 nm, Fig 2D) between the eGFP (*green*) and the PFR2 (*red*) signals along most of the cilium’s length (Figure 2D), showing that the detached cilium retained its attachment to the paraflagellar rod. We also found that the eGFP signal extended beyond the PFR2 signal near the tip (Figure 2D, inset). These observations suggest that TbLC2 localized to the axoneme, rather than the paraflagellar rod, as expected [29].

We further confirmed the subcellular localization of TbLC2 to the cilium using cell fractionation and immunoblotting. We sheared the cilia off ΔFLAM3/Rescued cells and separated them into cell body and ciliary fractions. We then prepared lysates from the whole-cell (WC), cell body only (CB), and ciliary (cilia) fractions and probed them by western blot using an anti-GFP antibody (Materials and Methods). We observed strong bands at approximately 50 kDa, which corresponds to the molecular weight of TbLC2::eGFP (Materials and Methods), in all the fractions of the induced cells, including the ciliary fraction (S4 Fig). This result confirmed the localization of TbLC2 to the cilium.

### TbLC2 knockdown causes morphological phenotypes associated with incomplete cytokinesis

We examined the effect of LC2 knockdown on the morphology of cultured trypanosome cells using wide-field microscopy (Materials and Methods). We found that ΔFLAM3/ΔLC2 cells formed clusters consisting of two to over ten cells after 72 hours of RNAi induction (Fig 3A, *top*), while uninduced and ΔFLAM3 cells did not (Fig 3A, *top*). We also found that ΔFLAM3/Rescued cells exhibited less clustering than ΔFLAM3/ΔLC2 cells (Fig 3A, *top*). These observations suggest that LC2 knockdown leads to a cell division phenotype.

**Fig 3.**
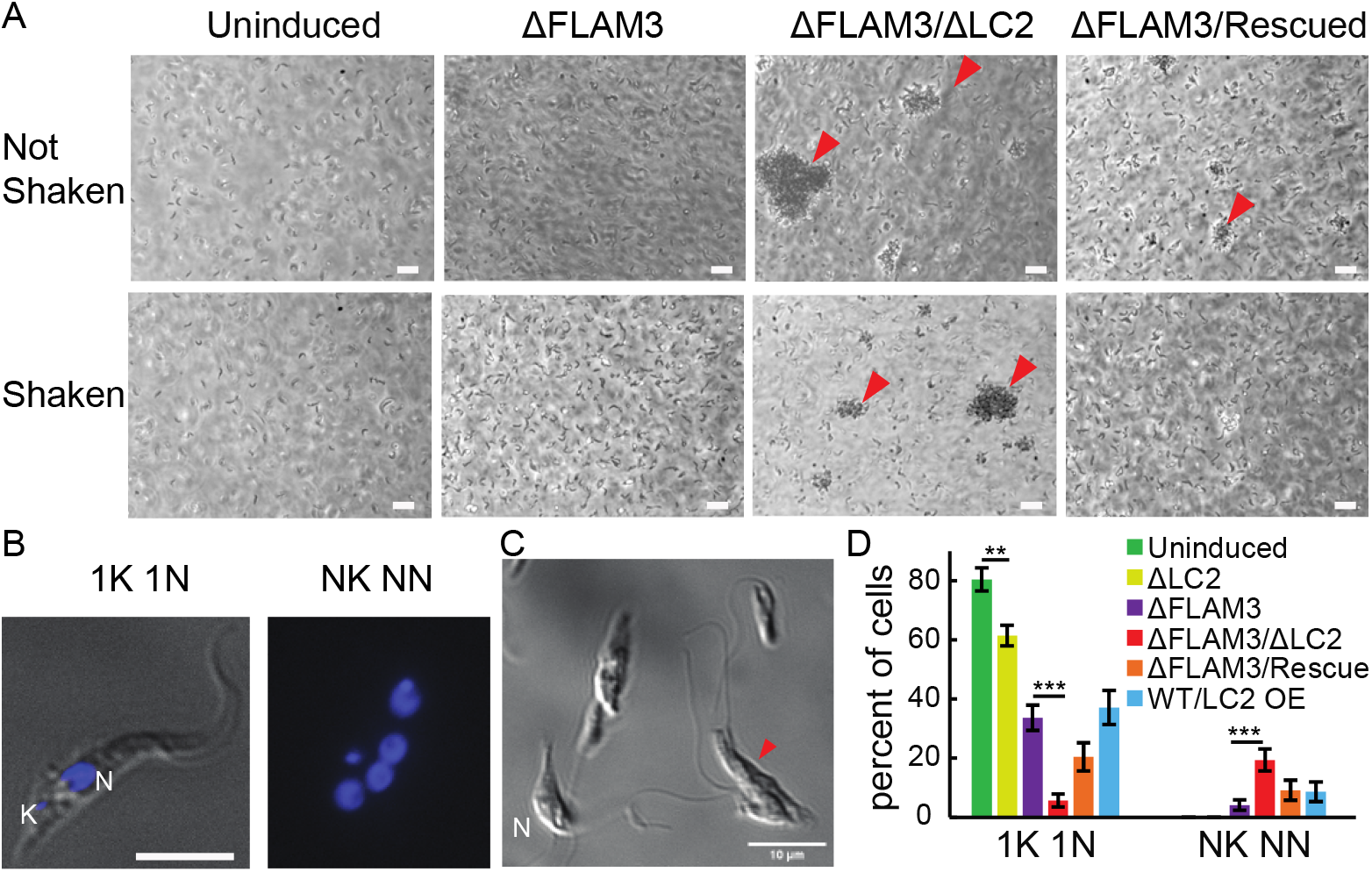
LC2 and FLAM3/LC2 knockdown cause mislocalization of kinetoplast and cell division defects. **A.** Representative images of uninduced, ΔFLAM3, ΔFLAM3/ΔLC2, and ΔFLAM3/Rescued cells cultures in the culture flask using phase-contrast microscopy 72 hours post-induction when we did not shake (*top*) and shook (*bottom*) the flasks. Major clusters of cells are indicated (*red arrows*). The scale bars represent 10 µm. **B.** Representative DAPI stained images for classification of kinetoplast (K) and nucleus (N) mislocalization. 1K 1N refers to normal cells with one kinetoplast normally localized to one nucleus (*left*). NK NN refers to cells classified as having >2 mislocalized (closer to each other) kinetoplasts and nuclei (*right*). The scale bar represents 5 µm and both images in this panel have the same scale. **C.** DIC image of fixed ΔFLAM3/ΔLC2 cells showing a representative amorphous clump of cells with multiple detached cilia (*red arrow*). **D.** Occurrence frequency of the normal kinetoplast and nucleus localization (1K 1N) and the most severe mislocalization (NK NN) phenotypes, as described in panel **B.**, in uninduced and induced (72 hours post-induction) ΔLC2, ΔFLAM3, ΔFLAM3/ΔLC2, ΔFLAM3/Rescued, and WT/LC2 OE cells. N=101, 192, 122, 111, 72, and 70 total classified cells of each strain, respectively. Other, less severe phenotypes, e.g., 1K 2N, 2K 1N, and 2K 2N, account for the percentages not represented. The error bars represent the statistical counting error. ** = *p*-value < 0.001 and *** = *p*-value < 0.00001, two-tailed paired t-tests.

Several *T. brucei* cell lines with cytokinesis-based cell division defects exhibit correlated motility phenotypes, including PFR2 [33, 34], dynein intermediate chain 138 (IC138) [35], dynein light chain 1 (LC1) [36], and trypanin [37] knockdown cells, and LC2 depletion causes ciliary motility defects [38, 39] in *C. reinhardtii*. Because we found that the TbLC2 knockdown cell division phenotype was particularly severe in FLAM3 knockdown cells, which exhibit altered ciliary motility due to the cilium being detached from the cell body (Fig 3A), we hypothesized that cell motility-like fluid shear forces induced by shaking the cell cultures might rescue TbLC2 knockdown cilium motility-based cell division phenotypes. We compared the propensity of the cells to form clusters in various cell lines when cultured with and without shaking (Materials and Methods). We found that gentle shaking (80-90 rpm) resulted in fewer and smaller cell clusters than no shaking in ΔFLAM3/ΔLC2 and essentially eliminated them in ΔFLAM3/Rescue cells (Fig 3A, *bottom*). These observations suggest that the TbLC2 knockdown cell division phenotype may be due, in part, to a motile cilium-based cytokinesis defect.

Replication of its cilium is an essential part of *T. brucei*’s cell cycle [40]. Basal body duplication, followed by nuclear and kinetoplast DNA replication, initiates cilium replication [41]. Therefore, abnormalities in cell cycle progression frequently result in nucleus and kinetoplast number and localization defects characterized by kinetoplasts being close to or sometimes indistinguishable from the nuclei [35]. We imaged DAPI-stained trypanosome cells to characterize the number and relative separation of the nuclei and kinetoplasts in multiple LC2 knockdown and control cell lines (Fig 3B, Materials and Methods). We found that LC2 knockdown cells were more likely to have mislocalized nuclei and kinetoplasts (representative images in Fig 3B), and ΔFLAM3/ΔLC2 cells tended to form amorphous clumps with multiple detached cilia (Fig 3C). By categorizing these phenotypes, we found the percentage of normal cells, i.e., with one nucleus well separated from one kinetoplast (1N 1K, Fig 3B), was 1.3-fold reduced in ΔLC2 and 2.3-fold reduced in ΔFLAM3 cells (*p-v*alues < 0.001 and < 0.00001, respectively, two-tailed paired *t*-tests, Fig 3D), as compared to uninduced cells. The mislocalization phenotype due to the LC2 knockdown was greatly exacerbated in the FLAM3 knockdown cells, with ΔFLAM3/ΔLC2 cells showing a 13.5-fold reduction in the fraction of cells with normal localization (*p*-value < 0.0001, two-tailed paired *t*-tests, Fig 3D), as compared to uninduced cells. Moreover, we observed that the most severe kinetoplast and nucleus mislocalization phenotype (NK NN, Fig 3B), which correlates to multi-ciliated amorphous cellular clumps (Fig 3C), occurred at significantly higher frequency in the ΔFLAM3/ΔLC2 than in ΔFLAM3 cells (4.7-fold, *p*-value < 0.00001, two-tailed paired *t*-tests, Fig 3D).

Cytokinesis defects in *T. brucei* often correlate either to defects in the length of the cilium or to the biophysical properties of the ciliary beat [37, 42], we compared the lengths of wild type cilia extracted using biochemical cell fractionation (Materials and Methods) and ΔFLAM3 and ΔFLAM3/ΔLC2 cilia extracted using mechanical shearing (Materials and Methods). We found that the wild-type cilia were 18.25 ± 0.42 μm (mean ± SEM, N = 76), ΔFLAM3 cilia were 16.80 ± 0.42 μm (N = 76), and ΔFLAM3/ΔLC2 cilia were 17.07 ± 0.34 μm (N = 52) long (S5 Fig). ΔFLAM3 and ΔFLAM3/ΔLC2 cells had slightly (< 10%) but significantly (*p-v*alues = 0.007 and 0.02, respectively, two-tailed paired *t*-tests) reduced ciliary length as compared to wild-type cells (S5 Fig). However, the length of ΔFLAM3 and ΔFLAM3/ΔLC2 cilia were not significantly different from each other (*p-v*alue = 0.64, two-tailed paired *t*-tests, S5 Fig), suggesting TbLC2 knockdown does not affect the length of the cilium.

Together, these observations demonstrate that knocking down TbLC2 disrupts cell division at cytokinesis after undergoing ciliary neogenesis. Because gentle shaking of the cell cultures partially rescues this phenotype, these observations further suggest that the TbLC2 knockdown-related cytokinesis phenotype may be due to ciliary motility defects. However, TbLC2 knockdown does not affect the length of the cilium, suggesting that TbLC2 may regulate the beating waveform of trypanosome cilia. This interpretation is consistent with the formation of trypanosome cell clusters due to abnormal cytokinesis in dynein light chain 1 (LC1, a regulator of ciliary beating) knockdown bloodstream and procyclic form *T. brucei* cells [36, 43], for example.

### TbLC2 is not essential for stable assembly of outer arm dynein in the axoneme

To examine whether the TbLC2 knockdown-related cell division and any motility phenotypes could arise from a difference in the stable assembly of axonemal dynein into the cilium, we visualized ciliary cross-sections with transmission electron microscopy (Materials and Methods). We found that both inner and outer arm dyneins were intact in the axoneme of ΔFLAM3/ΔLC2 cells (Fig 4, *middle*), like wild-type (Fig 4, *left*) and ΔFLAM3/Rescued (Figure 4, right) cells. This observation is unlike the *C. reinhardtii* Tctex-type LC2 outer arm dynein light chain mutant, in which outer arm dynein fails to assemble into the axoneme [38]. However, the observation is similar to other Tctex-type dynein light chain mutant cell lines, including Tcte3 knockout [21], a Tctex2 dynein light chain in mouse outer dynein arm, and Tctex2b knockout [44], a Tctex2-related inner dynein arm light chain in *C. reinhardtii*, which are not necessary for the stable assembly of their respective axonemal dynein arms into the axoneme. Other major structural components of the axoneme, including central pair and microtubule doublets, also appeared unperturbed by the lack of TbLC2 in the knockdown cells (Fig 4). Additionally, the cilium retained the paraflagellar rod (Fig 4), commensurate with the PFR2 immunofluorescence imaging (Fig 2D). Because the knockdown of TbLC2 did not prevent the assembly of outer arm dyneins into the axoneme (Fig 4) nor change the length of the cilium (S5 Fig) and yet resulted in cell division and kinetoplast localization defects associated with impaired ciliary motility (Fig 3), these results suggest that TbLC2 has a regulatory role in outer arm dynein function.

**Fig 4.**
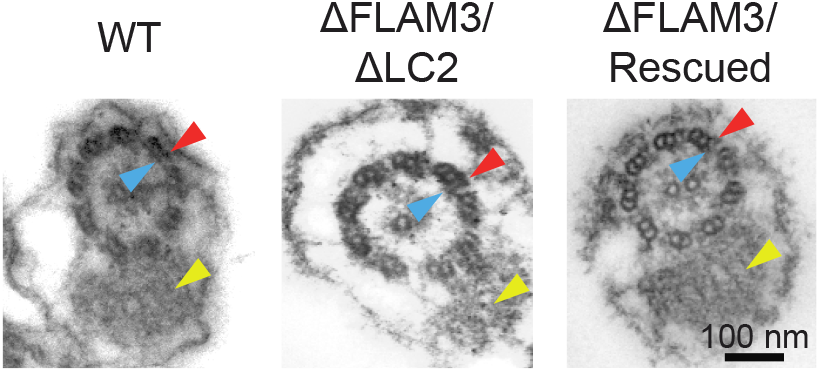
Stable assembly of outer arm dynein into the axoneme does not require TbLC2. Transmission electron microscopy (TEM) images of wild-type (WT, *left*), ΔFLAM3/ΔLC2 (*middle*), and ΔFLAM3/Rescued (*right*) cells show the axoneme (canonical 9+2 microtubule arrangement) and paraflagellar rod (*yellow arrowheads*). The outer arm (*red arrowheads*) and inner arm (*blue arrowheads*) dyneins were intact in all three cell lines. The scale bar is 100 nm, and all micrographs have the same magnification.

### LC2 knockdown results in reduced growth rates

To further quantify the effect of TbLC2 on *T. brucei* cell division, we examined the effect of the RNAi knockdowns on cell growth rate (Materials and Methods). The ΔFLAM3/ΔLC2 cells grew at a significantly reduced rate, with a population doubling time of 18.0 ± 1.4 hours, as compared to 14.5 ± 0.2 hours for ΔFLAM3 cells (*p-v*alue < 0.05, two-tailed paired *t*-test, doubling time determined in the first 72 hours of culture, Fig 5A). We also found that ΔFLAM3/ΔLC2 cells grew to approximately 1/10^th^ and 1/6^th^ the density of the uninduced and ΔFLAM3 cells after six days post-induction (*p-v*alues < 0.0001 in both cases, two-tailed paired *t*-tests, Fig 5A), respectively. These results are consistent with the growth defects reported in the case of other ciliary protein knockdowns, including LC1 [36], a leucine-rich outer arm dynein light chain, and Centrin3 [45], an inner arm dynein IAD5-1 associated protein in *T. brucei*. The results provide further evidence that TbLC2 knockdown-based motility defects lead to incomplete cytokinesis during cell division. The growth phenotype was rescued, in part, by overexpression of TbLC2::eGFP in ΔFLAM3/Rescued cells, as shown by a small (1.6-fold) but significant (*p-v*alue = 0.012, two-tailed paired *t*-test) increase in the cell density as compared to ΔFLAM3/ΔLC2 cells after six days (Fig 5A).

**Fig 5.**
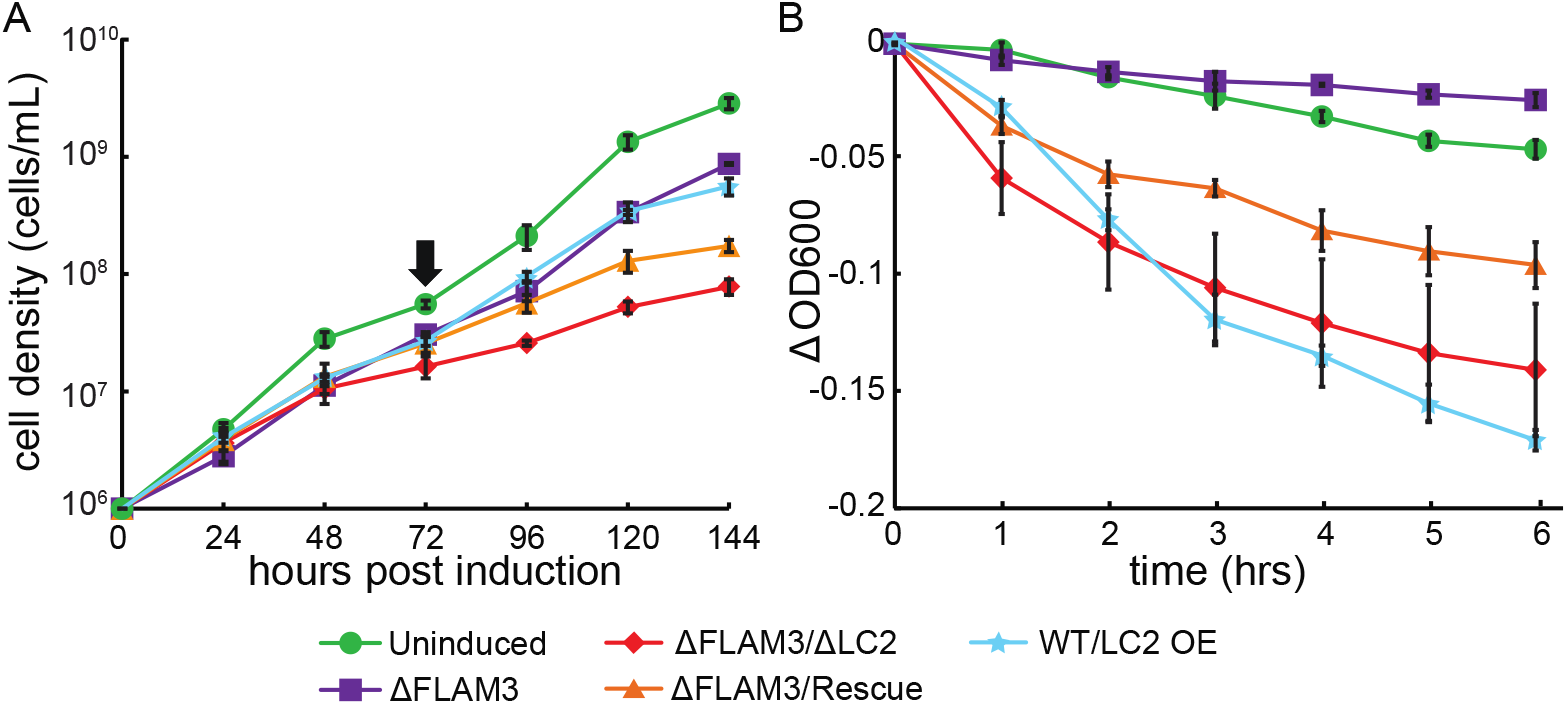
LC2 knocked down cells show growth and motility defects. **A.** Growth curves of various RNAi knocked down cells and uninduced cells (*green*). At 72 hours post-induction (*black arrow*), we diluted the cells back to the starting cell density (1×10^6^ cells/mL) and allowed them to grow for another 72 hours. The cell densities after 72 hours are scaled back from the diluted culture volume to reflect the total growth. **B.** Sedimentation curves of various RNAi knocked down cells and uninduced cells (*green*). In both panels, each data point represents the mean from three independent experiments, and the error bars represent the standard error of the mean (SEM). The legend applies to both panels.

Moreover, we found that the ΔFLAM3/ΔLC2 cells grew to the approximately same density as ΔFLAM3 cells during the first 48 hours post-induction (*p-v*alue = 0.81, two-tailed paired *t*-test, Fig 5A). We only found a significant reduction in growth after 48 hours post-induction (1.9-fold reduced cell density at 72 hours, *p-v*alue = 0.005, two-tailed paired *t*-test, Fig 5A) and beyond the first passage of the cells at 72 hours post-induction (2.1-fold reduced doubling rate, *p-v*alue < 0.001, two-tailed paired *t*-test, Fig 5A). We attributed the higher cell culture growth rates during the initial days after RNAi knockdown to the endogenous expression level of the proteins initially present at induction, the long-lived nature of proteins assembled into axonemes, and that the cell cultures grow slower only once the population of cells with cilia built during cell cycles that occurred after induction becomes significant.

Additionally, to investigate if the observed reduction in growth rate of the ΔFLAM3/ΔLC2 cells (Fig 5A) is the direct result of cells’ inability to segregate due to inadequate shear force generated by ciliary beating, we monitored the growth of ΔFLAM3/ΔLC2 cells while continuously maintaining the culture flask on an orbital shaker at 80-90 rpm (Materials and Methods). We found no significant increase in growth rate between the culture flasks with and without shaking (*p-v*alue > 0.05, two-tailed paired *t*-tests, S6 Fig). This result suggests an indirect effect of TbLC2 knockdown on cytokinesis that external shear forces could not rescue to induce cell separation [37].

### LC2 knockdown results in a rapid settling of the cells in solution

To investigate the effect of TbLC2 depletion from wild-type and FLAM3 knockdown cells on their swimming behavior, we performed a sedimentation assay by monitoring the change in absorbance near the top of cell solution at 600 nm over time (Materials and Methods). We found that ΔFLAM3/ΔLC2 cells settled to the bottom of the cuvette at a faster rate than the uninduced or ΔFLAM3 cells, as determined by the change in OD600 after 6 hours (*p-v*alues = 0.009 and 0.004, two-tailed paired *t*-tests, Fig 5A) respectively, Fig 5B), and as was previously reported in the case of silencing of several other ciliary proteins [36,38,45]. This motility phenotype was rescued, in part, by overexpression of TbLC2::eGFP in ΔFLAM3/Rescued cells, as shown by change in the OD600 value that was not statistically different from ΔFLAM3/ΔLC2 cells (*p-v*alue < 0.05 at 6 hours, two-tailed paired *t*-tests, Fig 5B). Together, these results suggest that TbLC2 is necessary for the effective swimming motility of trypanosome cells.

### Overexpression of TbLC2 only partially rescues cell growth and motility defects

The overexpression of recombinant TbLC2::eGFP in ΔFLAM3/Rescued cells only partially rescued the cell division (Figure 3), growth (1.4-fold increase after six days post-induction, Fig 5A), and sedimentation (1.5-fold decrease in ΔOD600 after 6 hours, Fig 5B) phenotypes associated with TbLC2 knockdown. We hypothesized that the partial rescue was due to overexpression of TbLC2::eGFP, rather than rescuing to the endogenous expression level. To test this hypothesis, we examined the phenotypes associated with overexpression of TbLC2::eGFP in wild-type cells (WT/LC2 OE cells), which express the endogenous TbLC2 protein as well. We compared the expression level of the recombinant LC2::eGFP in the WT/LC2 OE and ΔFLAM3/Rescued cells using flow cytometry (Materials and Methods) and found a similar level of eGFP fluorescence, with both cell lines showing a higher fluorescence intensity than the background fluorescence signal shown by the WT cells (S3 Fig).

The WT/LC2 OE cell line showed a 2.2-fold decrease in the fraction of normal 1K 1N cells as compared to wild-type (*p-v*alue <0.0001, two-tailed paired *t*-test, Fig 3D), and it showed a non-zero fraction of the cells exhibiting the most severe kinetoplast localization (NK NN, *p-v*alue = 0.003, two-tailed *t*-test, Fig 3D), as opposed to uninduced cells, which did not exhibit the NK NN phenotype at all (Fig 3D). Additionally, the WT/LC2 OE cell line showed a significant growth defect, with a 2.6-fold decrease in cell density at the end of 6 days (*p-v*alue < 0.001, two-tailed paired *t*-test, Fig 5A), as compared to wild-type cells. Moreover, WT/LC2 OE cells showed a significantly faster sedimentation rate (*p-v*alue < 0.001 at 6 hours, two-tailed paired *t*-test, Fig 5B) than uninduced cells. Together, these results show that overexpression of TbLC2::eGFP is sufficient to cause significant growth and motility defects. When taken with the growth and motility defects observed in the TbLC2 knockdown cell lines, these results suggest that there may be an optimal expression level of TbLC2 in trypanosome cells. They suggest how the overexpressed nature of recombinant TbLC2::eGFP in the ΔFLAM3/Rescued cells could cause the partial rescues of the growth and motility phenotypes observed in the ΔFLAM3/ΔLC2 cells back to the wild-type phenotypes despite efficient incorporation of TbLC2::eGFP into the cilium (Fig 2D and S4). However, whether the mechanism behind this discrepancy is due to changes in cell physiology caused by excessive TbLC2::eGFP, resource strain placed on the cells to produce the excess TbLC2::eGFP, the presence of the BCCP, eGFP, or His6 tags on the recombinant LC2, or some other factor, remains unknown.

### LC2 knockdown upregulates ciliary beat frequency

The length, beat frequency, and beating waveform are the primary biophysical properties of cilia that determine the swimming motility of ciliated microorganisms [12]. The lengths of ΔFLAM3 and ΔFLAM3/ΔLC2 cilia were not significantly different (*p*-value = 0.32, two-tailed paired *t*-test, S5 Fig), suggesting that length cannot explain the effect of TbLC2 knockdown on the swimming motility phenotype. Therefore, we examined the beat frequency of the various TbLC2 knockdown strains. To do so, we used a single-beam optical tweezer because of its ability to get high resolution frequency data with cells that are minimally constrained (Fig 6A, Materials and Methods). We found that the cells trapped at a single point displayed unconstrained ciliary beating behavior, with the cells performing a non-planar ciliary beat, like freely swimming cells (S1 Movies). We also found that the laser trapped both wild-type and FLAM3 knockdown cells, which have different body shapes and ciliary attachments (Fig 2), at a point approximately 2/3 of the cell length toward the posterior end (Fig 6B and S1 Movies). The specific location of the trapped position on the cells towards the posterior end suggests the cells have a localized structure suitable for optical trapping. Since the relative refractive index of the trapped material determines the intensity of gradient force exerted by the laser [46], DNA (mitochondrial, nuclear, or in the kinetoplast, in *T. brucei*’s case), which has a relatively high refractive index, in the cell body could determine the preferred trapping location [47].

**Fig 6.**
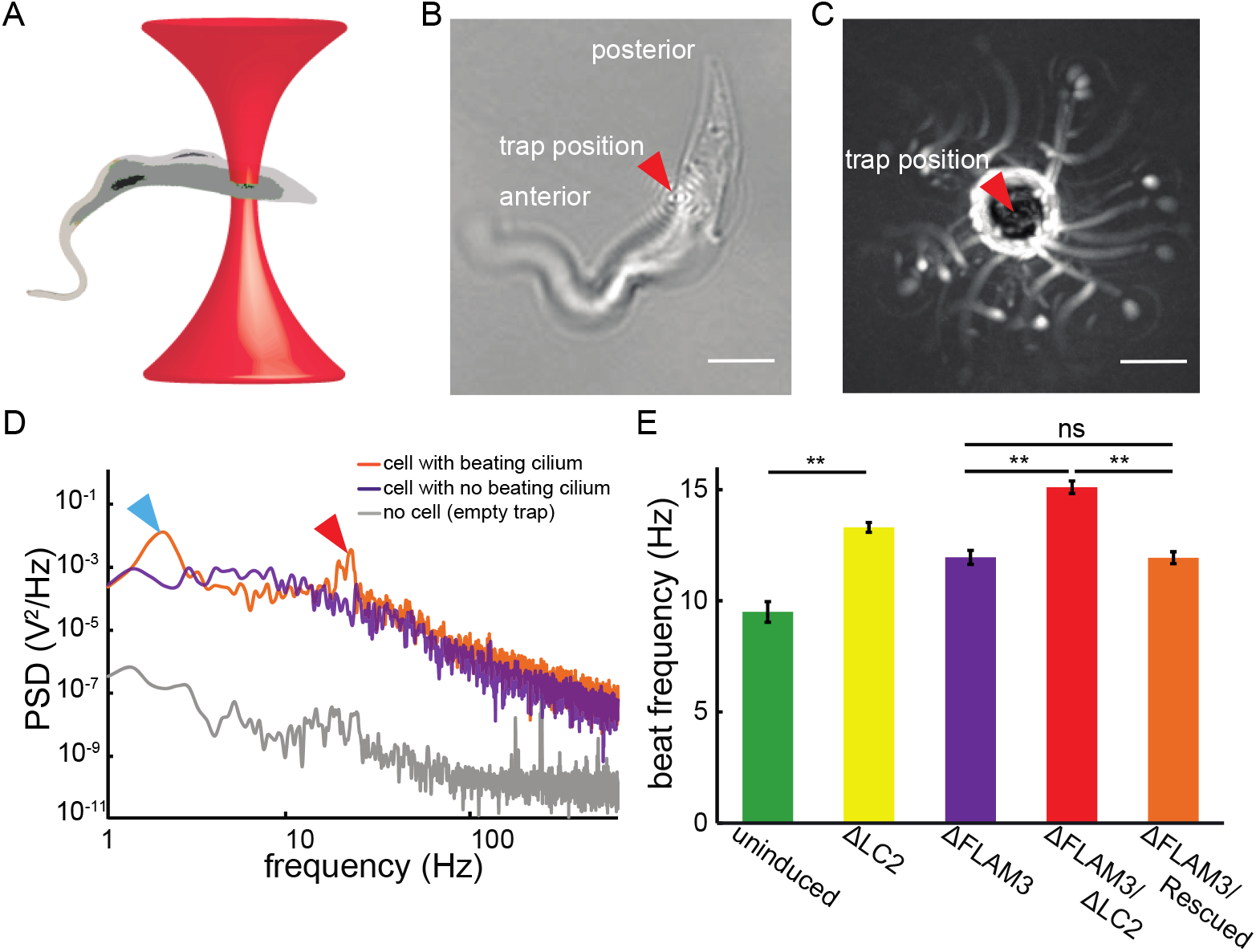
Optically trapped ΔLC2 and ΔFLAM3/ΔLC2 cells showed upregulated ciliary beat frequency. **A.** Schematic of a *T. brucei* cell (*gray*) trapped by a low power (∼20 mW) optical tweezer (*red*). **B.** Brightfield image of an uninduced ΔFLAM3/ΔLC2 cell showing the typical trap location on a trypanosome cell (*red arrowhead*). Scale bar = 5 µm. **C.** Typical maximum projection (intensity inverted) of a movie (2 s) showing a trapped ΔFLAM3 cell as it rotates about the trap center (*red arrowhead*). Scale bar = 5 µm. **D.** Typical power spectral densities (PSD) of a trapped cell with its cilium beating (ΔFLAM3/Rescued cell, *orange*), a trapped cell with its cilium not beating (ΔFLAM3/Rescued cell stalled the bottom of the imaging chamber, *purple*), and background trap noise (*gray*, about six orders of magnitude smaller than the PSD of the cell with a beating cilium at the beat frequency). Peaks in the PSD represent the characteristic cell rotation rate (*blue arrowhead*) and ciliary beat frequency (*red arrowhead*). Each PSD represents the average of 3 spectra (1 second of data, each). **E.** Ciliary beat frequency, *f_ω_*, of multiple cell lines obtained from the higher of the two characteristic frequencies from the PSD analysis. The error bars represent the SEM and ** represents *p*-values < 0.00001 and ns represents p-values > 0.05 from two-tailed paired *t*-tests. N=25, 86, 75, 85, and 96 for uninduced, ΔLC2, ΔFLAM3, ΔFLAM3/ΔLC2, and ΔFLAM3/Rescued cells, respectively.

We found that trypanosome cells trapped by the laser continue to swim and that the trap causes them to rotate about the center of the trap (Fig 6C). We further analyzed their swimming using power spectral density (PSD) analysis (Materials and Methods). We found that the typical PSD had two peak frequencies: *f_0_* at approximately 1-2 Hz and *f_ω_* at approximately 10-15 Hz (Fig 6D). The lower frequency, *f_0_*, represents the rotational motion of the cell swimming about the trap location, a motility mode that arises from the moment generated by the ciliary propulsive force acting about the trap center. We corroborated this interpretation of *f_0_* using maximum projections of time-series image sequences, which showed cells rotating about the trap location at this frequency (Fig 6C).

Due to the hydrodynamic forces acting on a free-swimming trypanosome cell with a cilium that wraps around the cell body, i.e., extensively in wild-type cells but only minimally in FLAM3 knockdown trypanosome cells, forward swimming motility generates a rotation of the cell about its long body axis at a frequency of approximately 2 Hz (5-8 single ciliary beats per full rotation) [48]. However, with no net forward propulsion of cells under the optically trapped condition, the cells do not rotate about their long body axis [49, 50], and we did not see a third peak in the PSDs of beating cells at about 2 Hz (Fig 6D). Therefore, we found that the higher frequency, *f_ω_*, corresponded to ciliary beat frequency.

We found that the beat frequencies, *f_ω_* from PSDs, of both ΔLC2 (13.3 ± 0.2 Hz, mean ± SEM) and ΔFLAM3/ΔLC2 (15.1 ± 0.3 Hz) cells were significantly higher than the beat frequencies of uninduced (9.5 ± 0.46 Hz) and ΔFLAM3 (11.95 ± 0.31 Hz) cells, respectively (*p*-values < 0.0001 in both cases, two-tailed paired *t*-tests Fig 6E). Overexpressing TbLC2::eGFP in the ΔFLAM3/Rescued cells restored the ciliary beat frequency to the parent FLAM3 knockdown ciliary beat frequency (11.9 ± 0.3 Hz, *p*-value = 0.49, two-tailed paired *t*-test Fig 6E). Together, these results show that knockdown of TbLC2 upregulates the beat frequency and thus suggests that TbLC2 downregulates the ciliary beat frequency in wild-type trypanosome cells, which is consistent with the regulatory role of other axonemal dynein associated proteins in *C. reinhardtii* [12] and *L. mexicana* [19].

### Ciliary motility is less directional in TbLC2 knockdown cells than wild-type cells

The combination of an elevated ciliary beat frequency (Fig 6E), which indicated knocking down TbLC2 upregulates motility, with an increased sedimentation rate (Fig 5B), which indicated knocking down TbLC2 downregulates motility, suggests a more complicated role for LC2 in regulating trypanosome cell swimming behavior, potentially involving the directional persistence of the swimming path. To investigate the role of TbLC2 in the directional persistence of trypanosome cell swimming paths, we recorded movies of freely swimming trypanosome cells (Materials and Methods, S2 Movies for examples). We analyzed the movies for individual cell swimming trajectories (Materials and Methods) and plotted the 10-second trajectories from a single origin for various cell strains (Fig 7A). The longer swimming trajectories suggest that uninduced, ΔLC2, and ΔFLAM3 cells exhibited more directionally persistent motility than ΔFLAM3/ΔLC2, ΔFLAM3/Rescue, and WT/LC2 OE cells (Figure 7A). Most strikingly, we found that ΔFLAM3/ΔLC2 cells exhibited less directionally persistent phenotype than ΔFLAM3 cells (Fig 7A).

**Fig 7.**
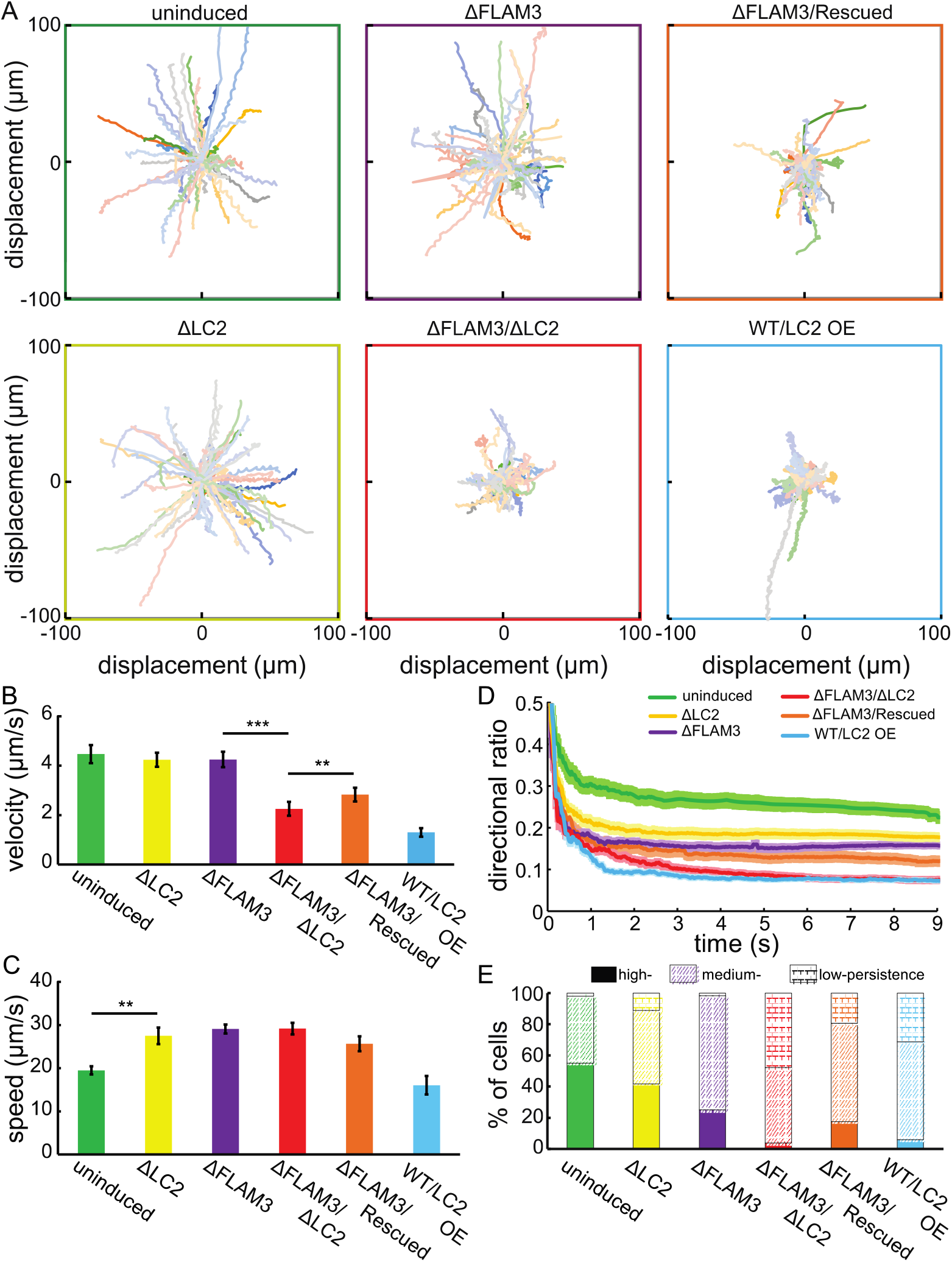
LC2 knockdown trypanosome cells have non-processive swimming behaviors. **A.** Swimming trajectories of trypanosome cells plotted from an initial position at the origin. Each trajectory represents 10 seconds of tracked data. **B.** Average velocity of swimming trypanosome cells. **C.** Average cell curvilinear swimming speed measured for multiple cell lines over the entire trajectory of 10 s. Error bars represent the SE of the mean, *** corresponds to *p*-values < 0.00001, and ** corresponds to *p*- values < 0.003, two-tailed paired *t*-tests, in panels A. and B. **D.** The mean directional ratio (DR, dark lines) calculated over elapsed time and averaged for each strain. Error bars (lightly shaded bands) represent the SE of the mean. **E.** The percentage of cells that exhibit high persistence (DR > 0.2, solid color), medium persistence (0.05 < DR < 0.2, hatch mark), and low persistence (DR < 0.05, brick pattern), as determined by the last point (as time goes to infinity) directional ratio. We showed paths for and calculated values from N = 50, 75, 53, 73, 47, and 68 for uninduced, ΔLC2, ΔFLAM3, ΔFLAM3/ΔLC2, ΔFLAM3/Rescued, and WT/LC2 OE cells, respectively, in all panels.

A closer look into the video microscopy (S2 Movies) showed that the shorter and less directionally persistent swimming trajectory phenotype associated with TbLC2 knockdown was characterized by cells that either mostly tumbled or swam short distances between tumbling events, as compared to the uninduced or ΔFLAM3 cells. We also found that the ΔLC2 and ΔFLAM3/ΔLC2 cells maintained a vigorous ciliary beat frequency (S2 Movies and Fig 6E), despite the propensity to tumble. Together, these observations suggest a role for TbLC2 in regulating the directionality of ciliary-driven swimming motility in trypanosomes.

To quantify the differences in directional persistence of the trypanosome cells (Fig 7A), we calculated the mean of the magnitude of the average velocity, which is the rate of the displacement of the cell between the start and final point of the motility track (Materials and Methods). We found that uninduced (4.62 ± 0.37 µm/s, mean ± SE of the mean) and ΔFLAM3 (4.48 ± 0.30 µm/s) cells had similar average velocity (*p*-value = 0.77, two-tailed paired *t*-test, Fig 7B), as previously reported [51]. ΔLC2 cells (4.24 ± 0.20. µm/s) swam with slightly (1.1-fold), but not significantly (*p*-values = 0.14, two-tailed paired *t*-test, Fig 7B), reduced average velocity, in accordance with our observations of trajectory length (Fig 7A). The reduced average velocity phenotype was exacerbated in ΔFLAM3/ΔLC2 cells (1.79 ± 0.18 µm/s, 2.5- fold reduced average velocity as compared to ΔFLAM3, *p*-value < 0.0001, two-tailed paired *t*-test, Fig 7B).

We also calculated the curvilinear swimming speed, which characterizes how fast the cells move along their actual path (Materials and Methods). We found that the swimming speed was higher for the ΔLC2, ΔFLAM3, and ΔLC2/ΔFLAM3 cells (27.5 ± 1.93 µm/s, 29.1 ± 1.0 µm/s, 29.2 ± 1.4 µm/s, respectively), as compared to that of uninduced cells (19.5 ± 0.9 µm/s, *p*-values < 0.001 in all cases, two-tailed paired *t*-tests, Fig 7C). This speed was in good agreement with the WT speed as reported in numerous previous studies [18,51–53]. We also found that the faster curvilinear swimming speed phenotype was partially restored to the lower uninduced speed in the ΔFLAM3/Rescued cells (25.7 ± 1.7 µm/s, *p*-value = 0.001, two-tailed paired *t*-test, Fig 7C).

The partial rescue of curvilinear swimming speed and straight-line velocity in ΔFLAM3/Rescued cells could be attributed to the overexpression of TbLC2 in the rescue cell lines. WT/LC2 OE showed motility defects marked with a 3.5-fold (Fig 7B) and insignificant 1.2-fold reduction (Fig 7C) in the average velocity and average curvilinear speed (*p*-value < 0.001 and 0.08, two-tailed paired *t*-test, respectively) compared to uninduced cells. These observations reiterate the notion that overexpression of TbLC2::eGFP alone is sufficient to cause motility defects in the cells and that there may be an optimal TbLC2 expression level.

The faster curvilinear swimming speed of the LC2 knockdown cells (Fig 7C) precludes the possibility of significant defects in ciliary propulsive force generation explaining the reduced mobility (Fig 7A). Additionally, a lower average velocity than the curvilinear swimming speed in the LC2 knockdowns suggests a highly futile swimming behavior with low directional persistence. We used the directional ratio, DR, defined as the ratio of displacement to path length (Materials and Methods [54]), to quantify trypanosome cell motility directional persistence. In accordance with the degree of the discrepancy between the curvilinear swimming speed and average velocity measurements, both the ΔLC2 and ΔLC2/ΔFLAM3 cells suffered a loss in directional persistence along the entire trajectory with a lower directional ratio (DR = 0.18 ± 0.01 and 0.07 ± 0.01, respectively, mean ± SE of the mean, Fig 7D) at the final point of the swimming trajectory as compared to uninduced (0.23 ± 0.01, *p*-value = 0.004 and < 0.0001, respectively, two-tailed paired *t*-test) cells, with the double knockdown enduring the more severe defect of the two (Fig 7D).

Persistent ciliary motility is maintained by highly regulated forward (tip-to-base) ciliary beating. Disruption to such regulation or transient and abrupt switching of the ciliary beating mode leads to cellular reorientation (tumbling) [18]. To characterize the relative occurrence of tumbling in the population, we binned swimming cells into three directional persistence classes, high persistence (DR > 0.2, persistent swimmers, S2A Movie), medium persistence (0.05 < DR < 0.2, persistent swimmers with intermittent tumbling, S2B Movie), and low persistence (DR < 0.05, tumblers, S2C Movie), based on their directional ratio at the end of the 10-second tracks (Fig 7D). We found that approximately 50% of the uninduced cell population exhibited high persistence while very few exhibited low persistence (Fig 7E), which is consistent with previous reports [18]. The ΔFLAM3 cells showed a significant decreased in the fraction of high persistence swimmers to approximately 25% (*p*-value = 0.0018, two-tailed paired *t*-test, Fig 7E) without significantly changing the fraction of low persistence swimmers (*p*-value = 0.97, two-tailed paired *t*-test, Fig 7E), as compared to uninduced cells. However, both the ΔLC2 and ΔFLAM3/ΔLC2 cells showed a significant increase in the low persistence swimmer population to 11% (*p*-value = 0.043) and 48% (*p*- value < 0.0001, two-tailed paired *t*-test in both cases), and a corresponding decrease in the high persistence swimmer population to 42% (*p*-value = 0.11) and 3.8% (*p*-value = 0.0005, two-tailed paired *t*- test in both cases), as compared to uninduced and ΔFLAM3 cells, respectively (Fig 7E). Overexpression of TbLC2::eGFP in ΔFLAM3/Rescued cells partially, but not fully, rescued the phenotype. A significantly larger fraction of ΔFLAM3/Rescued cells were highly persistent swimmers (17.4%, *p*-value = 0.011, two-tailed paired *t*-test) and lower fraction were low persistence swimmers (19.6%, *p*-value = 0.0016, two-tailed paired *t*-test) as compared to the ΔFLAM3/ΔLC2 cells (Fig 7E). Complementing the growth and motility phenotypes results (Fig 5), a majority of the WT/LC2 OE cells showed medium or low persistence swimming behavior with a significant decrease in the high persistent swimmer proportion compared to the uninduced cells (5.9% from 55.1%, *p*-value < 0.001, two-tailed paired *t*-test, Fig 7E) showing that the overexpression of recombinant TbLC2 might have hindered the full restoration of motility phenotypes in rescue cells.

Together, these motile cell trajectory results suggest that TbLC2 regulates axonemal dynein in a way that reduces the tumble bias of trypanosome cells. They suggest that it stabilizes the switching mechanism between forward propulsion, with tip-to-base bending wave propagation along the cilium, and tumbling-like cell reorientation, with base-to-tip bending wave propagation. This behavior is reminiscent of mechanisms that alter the tumble bias in run-and-tumble swimming behavior [17, 55].

### LC2 and FLAM3 double knockdown alters the ciliary waveform of trypanosome cells

The combination of increased curvilinear swimming speed (Fig 7C) and beat frequency (Fig 6E) with low total displacement (Fig 7A), average velocity (Fig 7B), and directionality ratio (Fig 7D and Fig 7E) all suggest that the directionality of ciliary motility must be compromised in TbLC2 knockdown cells. We found that this was due, in part, to the increased likelihood of tumbling motility, which is often associated with switching between tip-to-base to base-to-tip ciliary bending wave propagation swimming modes in TbLC2 knockdown cells (S3 Movies) [2, 56]. However, the role of TbLC2 in maintaining a regular beating waveform could be a significant contributing factor, as well.

Wild-type *T. brucei* cells exhibit auger-like movement [2] driven by the beating of a single cilium that wraps around the cell body (Fig 2B). The out-of-plane nature of the cilium and its beating waveform makes it difficult to use conventional quantitative analysis techniques to extract meaningful information about the ciliary beat waveform on wild-type cells [17]. This issue can be mitigated when the cilium is well isolated from the rest of the body, like in *Leishmania* cells [56–58]. We used RNAi knockdown of FLAM3 (Figs 2B-D, Materials and Methods) to replicate this phenotype in trypanosome cells. We tracked individual cilia from multiple cells undergoing ciliary beat propagation (Materials and Methods) from ΔFLAM3 and ΔFLAM3/ΔLC2 cells. We first calculated the tangent angle (ψ) of the cilium as a function of normalized arc length along the cilium’s length, (s/L) (S7 Fig), and then calculated the normalized curvature (κ/L) by taking the derivative of the tangent angle with respect to normalized arc length (Materials and Methods) and plotted it as a function of normalized arc length (Fig 8A and Fig 8B). We found that the beat waveforms of both ΔFLAM3 and ΔFLAM3/ΔLC2 cells were asymmetric, i.e., the maximum positive curvature, 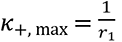, (Fig 8C) was greater than the maximum negative curvature, 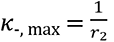, (Fig 8C), for a beat period. We calculated the maximum curvature asymmetry ratio, *AR*_MC,_ as the ratio of *κ*_+, max_ to *κ*_-, max_ (Materials and Methods), in both cases and found that *AR*_MC_ was larger in the ΔFLAM3/ΔLC2 cells (1.35 ± 0.03) than ΔFLAM3 cells (1.11 ± 0.06, *p*-value = 0.0006, two-tailed paired *t*-test, Fig 8D). Additionally, we found that the ΔFLAM3/ΔLC2 cells’ ciliary beat exhibited reduced peak-to- peak tangent angle amplitude (1.11 ± 0.03 radians, mean ± SEM, S7 Fig) as compared to ΔFLAM3 cells (1.25 ± 0.02, *p*-value = 0.004, two-tailed paired *t*-test, N=25 beating periods from approximately ten cells, S7 Fig). However, the beat wavelength as quantified by the normalized arc length between two consecutive tangent angle peaks along a beating cilium (S7 Fig) remained unaltered (0.96 ± 0.03 and 0.99 ± 0.05, *p*-value = 0.62, two-tailed paired *t*-test, N=25 beating periods from approximately ten cells) between the ΔFLAM3 and ΔFLAM3/ΔLC2 cell lines, respectively.

**Fig 8.**
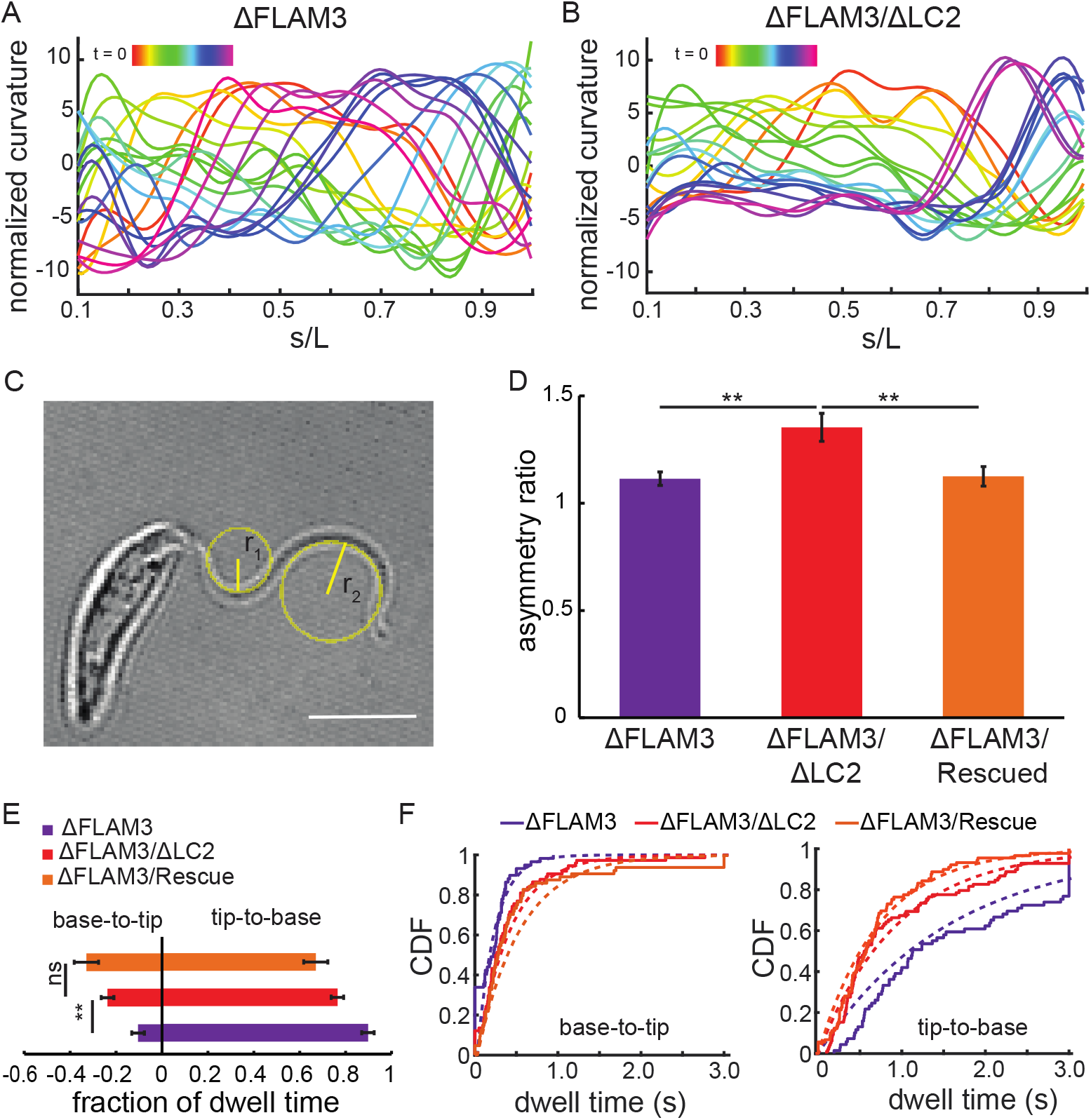
Loss of TbLC2 alters multiple aspects of the ciliary beat waveform. Representative plots of curvature, κ, normalized to the contour length (L) for **A.** ΔFLAM3 and **B.** ΔFLAM3/ΔLC2 cells plotted as a function of s/L, which is the ratio of the arc length along the cilium (s) to the total ciliary length (L), where s/L = 0 is the base and s/L=1 is the tip of the cilium. Colors (*red* to *magenta*) represent the time evolution of ciliary beat shape over one complete period of the oscillation. **C.** Schematic showing the radii of curvature corresponding to the maximum positive, r_1_, and the maximum negative, r_2_, curvature, where the curvature is 1/the radius of curvature, κ=1/r, in this frame. We used these radii of curvature to calculate the asymmetry ratio of a ciliary bend during a ciliary beating. This image is a representative frame from a high-speed movie of a ΔFLAM3/ΔLC2 cell. Scale bar = 5 μm. **D.** Mean maximum curvature asymmetry ratio, AR_MC_, for ΔFLAM3, ΔFLAM3/ΔLC2, and ΔFLAM3/Rescued cells. N = 15 waveforms from 10- 12 different cells in each cell line, the error bars represent the SEM, and ** represents a *p*-value < 0.005, two-tailed paired *t*-tests. **E.** Fraction of time cells dwelled in tip-to-base (positive fraction) and base-to-tip (negative fraction) ciliary beating modes. N = 42, 44, and 33 cells for ΔFLAM3, ΔFLAM3/ ΔLC2, and ΔFLAM3/Rescue cells, respectively. ** represents a *p*-value of < 0.001 and ns a *p*-value > 0.05, two-tailed paired *t*-tests. **F.** Cumulative distribution function (CDF) of individual dwell times for various FLAM3 knockdown cells in base-to-tip (*left*) and tip-to-base (*right*) ciliary beating modes. The fits (*dotted lines*) to exponential functions are shown. Forty-five cells of each strain were analyzed with a total number of n = 59, 75, 65 for base-to-tip mode and n = 70, 99, and 90 for tip-to-base mode for ΔFLAM3, ΔFLAM3/ ΔLC2, and ΔFLAM3/Rescue cells, switching events, respectively.

Cilia that exhibit a non-zero mean curvature (S7 Fig) in combination with a symmetric dynamic curvature (S7 Fig) propel straight-line swimming when the beats are synchronized in biflagellate cells, like *C. reinhardtii* [59], and leads to circular or spiral swimming trajectories in uniflagellate cells, like mammalian sperm [59]. Since *T. brucei* have a single cilium, effects that increase the asymmetry of either the static or forward swimming tip-to-base dynamic curvature could lead to curved swimming paths that reorient the cell body as it swims [59]. We also found that the intermittent base-to-tip ciliary beat mode of trypanosomes was highly asymmetric, with large maximum curvature bending wave propagation (S4 Movie and S8 Fig). Furthermore, ΔFLAM3/ΔLC2 cells showed a more highly asymmetric, non-zero static component of the ciliary beating waveform (S9 Fig) during a base-to-tip (reverse) ciliary beating than during the tip-to-base (forward) beating. Both of these observations correlated with the reorientation of the cell’s swimming direction upon reversal of its forward swimming tip-to-base beating to its reverse bast-to-tip mode (S3 Movies, Fig 7A, and Fig 7D).

To examine whether the LC2 knockdown shifted the biasedness of ciliary beating mode, we investigated the fraction of the time the cells dwelled in the reversed base-to-tip beating mode versus the forward tip-to-base mode (Materials and methods). We found that ΔFLAM3/ΔLC2 cells dwelled in the base-to-tip beating mode approximately twice as long as ΔFLAM3 knockdown cells (0.24 ± 0.027 and 0.10 ± 0.02, respectively, *p*-value < 0.001, two-tailed paired *t*-test, Fig 8E). We further analyzed these fractions by quantifying the duration (dwell time) of the individual ciliary beat mode events (Fig 8F). Exponential fits to the cumulative probability distribution functions of the dwell times revealed characteristic dwell times of 0.21 ± 0.03, 0.40 ± 0.04, and 0.52 ± 0.07 seconds for base-to-tip beat (reverse) mode and 1.57 ± 0.19, 0.95 ± 0.10, and 0.73 ± 0.08 seconds for tip-to-base (forward) mode (characteristic time of the exponential fit ± standard error of the fit in all cases) for ΔFLAM3, ΔFLAM3/ΔLC2, and ΔFLAM3/Rescue cells, respectively. The ΔFLAM3/ΔLC2 cells dwelled approximately 2-fold longer in each reverse base-to-tip event and 1.7-fold shorter in the forward tip-to-base beating mode events than ΔFLAM3 cells (*p*-value = 0.0025 and *p*-value < 0.0001, two-tailed paired t-tests, respectively). These results showed that the TbLC2 knockdown cells more rapidly switched into the base-to-tip (reverse) beating mode and were slower to switch back to the tip-to-base (forward) beating mode than those cells without TbCL2 knocked down. This result suggests that TbLC2 promotes directional persistence in trypanosome swimming paths by biasing cilium beating mode switching mechanism toward the forward swimming, tip-to-base beating mode.

Together, these observations and results suggest that TbLC2 regulates the fundamental beat frequency, the frequency at which the bending wave propagation direction switches, and the asymmetry in the static and dynamic components of the waveform of beating trypanosome cilia. With the asymmetric base-to-tip reverse ciliary beating directly related to cellular reorientation during ciliary motility, our data suggest that TbLC2’s regulation of beating mode directly controls the cell tumbling and ultimately the swimming directionality in *T. brucei*.

## Discussion

In this study, we identified a *Trypanosoma brucei* homolog of *Chlamydomonas reinhardtii* LC2 (Fig 1), a Tctex-type axonemal dynein light chain, that we named TbLC2 (Tb927.9.12820 [25]). We demonstrated that TbLC2 localizes to the axoneme (Fig 2) and regulates axonemal dynein-driven ciliary beating, swimming motility, and associated functions (Figs 3-8) through various molecular biological, cell biological, and cell biophysical techniques. Our results suggest that TbLC2 is not required for dynein assembly into the axoneme (Fig 4), unlike multiple other Tctex-type dynein light chains, e.g.,

*Chlamydomonas reinhardtii* LC2. We found TbLC2 knockdowns exhibit cell division and cytokinesis defects (Fig 3), reduced cell culture growth rates (Fig 5A), impaired cellular swimming motilities (Fig 5B), and lower directional swimming persistence (Fig 7). However, our results showed that TbLC2 knockdowns beat more rapidly. To reconcile this mechanism, we took detailed look at the beating waveforms of the TbLC2 knockdown cells and found that they beat more erratically than cells expressing wild-type levels of TbLC2. Specifically, we found that TbLC2 downregulates the ciliary beating frequency (Fig 6), reduces the magnitude of the static (time-averaged) curvature of beating trypanosome cilia (Fig 8), maintains dynamic ciliary curvature symmetry (Fig 8), and shifts the ciliary beat mode bias toward the canonical tip-to-base bending wave propagation direction-based highly persistent forward motility mode (Fig 8). Thus, we propose that TbLC2 keeps the ciliary beat of trypanosomes under control. Without the regulatory effects of TbLC2 on axonemal dyneins within the cilium, trypanosome cells are going nowhere fast.

### Lack of TbLC2 modulates ciliary waveform and correlates with less directional swimming behaviors

Persistent swimming is essential for directional migration through the tsetse fly vector, social motility [60, 61], and chemotaxis [62] within the highly viscous fluid and tissue environment of the trypanosome’s mammalian host. Swimming motility is a significant virulence factor [2,6,16] because it is necessary for trypanosome cells to cross the blood-brain barrier and to evade the immune system [5]. Trypanosomes achieve persistent directional swimming trajectories by maintaining their forward tip-to-base beating waveforms for extended periods. Periods of persistent swimming in wild-type trypanosome cells are interrupted by occasional pauses followed by a tumble [17]. The tumbling occurs when the ciliary beat waveform switches from a symmetric (zero static curvature) tip-to-base beating waveform into an asymmetric (large static curvature) base-to-tip ciliary beating waveform [57, 59]. Thus, defects that cause trypanosomes to increase the asymmetry of their beating modes, increase the likelihood of switching to the tumbling-associated base-to-tip ciliary beating mode, or increase the time they spend in the base-to-tip ciliary beating mode will affect trypanosome cell swimming motility by reducing the directional persistence of their swimming trajectories. We found that TbLC2 RNAi knockdowns tended to cause the trypanosome cilium to beat faster (higher frequency, Fig 6), more erratically (longer time in and more frequent switching to the base-to-tip beat mode, Fig 8E and Fig 8F), and with a significant asymmetry to the beat (Figs 8A-D). Together these results suggest that TbLC2 regulates axonemal dynein to maintain the propulsive, persistent tip-to-base ciliary beating mode.

Our suggestion that TbLC2 regulates dynein is generally consistent with studies on other axonemal dynein-associated proteins. For example, knockdown of [36] and mutations to [43] outer arm dynein-associated light chain (LC1) and intermediate chain (DNAI1) [63] both shifted the bias away from tip-to-base waveform propagation directed motility to sustained base-to-tip ciliary beating in *T. brucei*. These results suggest that outer arm dynein and its regulation may be particularly important for maintaining propulsive tip-to-base ciliary beating because both LC1 and DNAI1 directly associate with outer arm dynein. Moreover, the proximal/distal asymmetry in the molecular composition of the outer arm dynein docking complex regulates the beat propagation direction in *T. brucei* and *L. mexicana* [19]. The aggregate of these results suggests that TbLC2’s mechanism is to act as a potent, direct regulator of trypanosome axonemal dynein motility and likely outer arm dynein, specifically.

TbLC2’s downregulation of the ciliary beat frequency (Fig 6E), which is consistent with other outer arm dynein light chains and regulatory proteins, support of suggestion that TbLC2 regulates outer arm dynein, specifically. Deleting the LC4-like outer arm dynein light chain in *L. mexicana* resulted in a significant increase in ciliary beat frequency and a substantial increase in swimming speed and velocity [19]. Similarly, disruption of Tcte3-3 outer arm dynein light chain Tctex2 in mouse spermatozoa led to an increase in ciliary beat frequency [21]. However, we cannot rule out a role with inner arm dynein because loss of dynein regulatory complex protein DRC3, a component of the nexin dynein regulatory complex (N-DRC) subunit that interacts with inner arm dynein g [12] in *Chlamydomonas* cells, also elevated average ciliary beat frequency.

We also found that TbLC2 knockdowns result in a more asymmetric waveform (Fig 8B and Fig 8D). This result is additionally consistent with other axonemal dynein light chains and regulatory proteins. For example, the inner arm dynein-related DRC3 mutants have reduced amplitude ciliary beat waveforms Chlamydomonas cells [12]. Furthermore, outer arm dynein-associated protein LC1 regulates the mechanochemical cycles of outer arm dynein during ciliary beating in Chlamydomonas cells [64]. These pieces of evidence for ciliary waveform modulation by dynein-related structures, combined with our findings of an elevated ciliary beat frequency and altered beat symmetry observed in TbLC2 knockdown cells, strongly support a regulatory role for dynein-related light chains in maintaining an effective ciliary beating waveform.

Some knockouts, knockdowns, or mutations of axonemal dynein-related light chains, including LC1 in *Chlamydomonas* [64] and trypanosomes [36], and LC2 in *Chlamydomonas* [38], have shown contrasting motility phenotypes to the ones reported in this study, including a reduced ciliary beat frequency. However, the deletions in these cases led to failures in outer arm dynein assembly, which the knockdown of TbLC2 did not cause (Fig 4). We found no noticeable defects in the axonemal structures, including the outer and inner arm dyneins (Fig 4). Therefore, direct comparison to many other cell biophysical phenotypic studies of axonemal dynein-associated light chains is difficult. Our results suggest that TbLC2 is unnecessary for the stable assembly and activation of dyneins in the axoneme, the preassembly of axonemal dynein complexes in the cytoplasm, or the transport of dynein complexes to the axoneme.

### Ciliary beat asymmetry, rather than waveform propagation direction, underlies changes in trypanosome cell swimming direction

Beyond the cellular and molecular mechanisms of TbLC2, our results suggest that asymmetry of the beat, rather than the bending wave propagation direction, dominates cell reorientation during ciliary motility. We found that the tumbling motility mode and the resulting cellular reorientation are not inherent to the *symmetric* base-to-tip ciliary beating.

We found that the TbLC2::eGFP overexpressing wild-type, WT/LC2 OE cells, spent long periods of time in the base-to-tip ciliary beating mode (S5 Movie). However, they did not exhibit the high static curvature, asymmetric reverse ciliary beating characteristic of ΔFLAM3/ΔLC2 cells (S8 Fig and S4 Movie). Instead, they exhibited a symmetric, low static curvature, base-to-tip ciliary beating mode (S5 Movie). Thus, they swam with relative directional persistence, albeit backward. Unfortunately, we could not quantify these detailed comparisons because performing the same ciliary waveform tracking for WT/LC2 OE as we did for FLAM3 knockdown strains was impossible without the detached cilium.

Furthermore, a higher ciliary beat frequency (TbLC2 knockdown cells, Fig 6E) should increase the swimming speed and persistence of cells in the absence of other effects. However, a higher asymmetry ratio (TbLC2 knockdown cells, Fig 8D) should decrease the swimming speed and persistence of cells in the absence of other effects. Because we found that TbLC2 knockdown cells exhibit decreased directional motility persistence (Fig 7D and Fig 7E), these results further support the conclusion that asymmetry of the base-to-tip ciliary beat waveform dominates the tumbling mechanism, rather than the base-to-tip waveform propagation direction, itself. Moreover, the reverse base-to-tip ciliary beating mode in wild-type and TbC2 mutant trypanosomes is more asymmetric than the forward tip-to-base beating mode [57, 58] (Fig 8A, Fig 8B, as compared to S7 Fig and S8 Fig). The conclusion asymmetry of the base-to-tip ciliary beat waveform dominates the tumbling mechanism is also consistent with the observation that *Chlamydomonas* cells with only one cilium (e.g., with an ablated second cilium) exhibit an asymmetric beating waveform and tend to rotate with minimal swimming persistence [65]. Other *Trypanosomatidea* like *Leishmania major* and *Angomonas deanei* also show intermittent, highly asymmetric, breaststroke-like base-to-tip ciliary beating that causes cell reorientation and a significantly less efficient translational motility [57] than symmetric beating modes. Thus, we suggest a model of trypanosome directional motility that relies on a reversal of beating waveform propagation direction to switch the ciliary beat from a symmetric to an *asymmetric* waveform, which is the principal biophysical effect leading to the reorientation of the cell.

### TbLC2 regulates axonemal dynein through possible LC-IC interactions

Our bioinformatic analysis suggests that TbLC2 is the trypanosome homolog of *Chlamydomonas reinhardtii* LC2 (Fig 1 and S1 Fig). Assuming trypanosome dynein complexes share structural homology with Chlamydomonas [20] and *Tetrahymena thermophila* [66, 67] axonemes, we suggest that TbLC2 localizes to the outer arm dynein light chain tower, a considerable distance (15 nm) from outer arm dynein’s ATP hydrolysis site. Given this distance, our results suggest that TbLC2 regulates the dynein function through an intricate network of LC-LC and LC-IC interactions or perhaps with the next nearest neighboring dynein motor along the axoneme, based on its location in the dynein complex [20]. Furthermore, this analysis suggests that TbLC2 may directly interact with the HC-IC complex as a heterodimer with the trypanosome homolog of Chlamydomonas LC9 [20]. Additionally, a recent crosslinking experiment showed that Chlamydomonas LC9, in turn, associates with IC1 and IC2 and thus forming an LC-IC block [68] that influences the function of outer arm dynein in Chlamydomonas cilia through an interaction with the dynein heavy chains. In sum, these results from *C. reinhardtii* suggest that TbLC2, also being a Tctex-like light chain, might be an indispensable piece of IC-LC block architecture responsible for regulating the activity of outer arm axonemal dynein. However, despite the similarities in the sequences and modeled structures (Fig 1), we observed less severe motility phenotypes in TbLC2 knockdown cells compared to the Chlamydomonas LC2 mutants. Though these differences may arise from the lack of an outer arm dynein assembly role for TbLC2 (Fig 4), the morphological and physiological differences in the ciliary motility between the two organisms suggest that TbLC2 has a divergent dynein regulatory role, as compared to Chlamydomonas LC2.

### Growth and morphology morphological phenotypes of LC2 knockdown are not fully restored by mechanical agitation

The ΔFLAM3/ΔLC2 double knockdown cell cultures developed multiple-cell clusters and multi- ciliated cellular clumps (Fig 3). These observations are consistent with numerous previous studies relating ciliary motility defects to cell morphology and growth defects [2,35,37] as well as the inability to drive cell separation during late cytokinesis [35, 36]. We also found that kinetoplasts were closer to the nucleus than expected (1/3^rd^ or less than the average separation in ΔLC2 and nearly adjoining in ΔFLAM3/ΔLC2 cells, Fig 3B and Fig 3D) in TbLC2 knockdown cell lines, as compared to their respective parent cell lines. These discrepancies could result from, and provide supporting evidence for, a ciliary motility-related disturbance in the timing of kinetoplast division and segregation (unpublished data reported by Ralston et al. [37]), as was demonstrated by using a reaction force model [69]. However, we found that mechanical agitation of the ΔFLAM3/ΔLC2 cell culture failed to eliminate cell cluster formation (Fig 3A) and failed to improve cell growth (S6 Fig), suggesting that TbLC2 depleted cells may have additional physiological or morphological defects as a direct or indirect result of the motility defect that might have led to cytokinesis failure [37]. On the other hand, cluster and amorphous clump formation were reduced by shaking during ΔFLAM3/Rescued cell (Fig 3), signifying that rescuing the depletion of TbLC2 by overexpressing TbLC2::eGFP only partly restored the cytokinesis defect in the ΔFLAM3/ΔLC2 cells compared to its parent, ΔFLAM3, cell line (Fig 3).

### The effect of TbLC2 knockdown on cell morphology and motility is more pronounced in ΔFLAM3 than in wild-type parental cells

The quantitative analyses that we present of microscopic cell motility and cell growth showed significant phenotypes with TbLC2 RNAi knockdown in both wild-type (ΔLC2) and FLAM3 knockdown (ΔFLAM3/ΔLC2) cells (Fig 5 and Fig 7), with the effect being more severe on the latter cell line. In an attempt to understand how FLAM3 knockdown caused amplified TbLC2 phenotypes, we considered a recent analysis of trypanosome cell swimming directional persistence that showed morphologically straighter trypanosome cells followed straighter swimming trajectories than flexible cells, suggesting a direct correlation between cell stiffness and directionality in ciliary motility [18]. Additionally, results from *Spiroplasma*, a bacterium that does not have cell walls and swims in helical paths similar to *T. brucei*, also suggest that less rigid cells are more susceptible to fluctuations and perturbations, leading to reduced swimming directional persistence [49]. A subpellicular corset of microtubules arranged underneath the plasma membrane provides structural integrity and rigidity to the trypanosome’s cell body [70, 71]. When the cilium remains wrapped around the cell body with a mechanically rigid constraint, as the flagellar attachment zone provides in wild-type trypanosome cells, the cell body-subpellicular microtubule array- cilium complex acts in synergy and becomes stiffer (with larger effective flexural rigidity) than the arithmetic sum of the cilium and cell body stiffnesses. When the cilium becomes detached, it loses this synergy and becomes much softer.

Moreover, the chiral nature of the trypanosome cilium’s attachment to its cell body leads to an asymmetry in the hydrodynamic drag experienced by the cell during swimming motility, causing rotation of the trypanosomes about their long cell body axis, which is a dominant contributor to propulsive thrust and directionality of swimming [56]. Although the ciliary beat of FLAM3 knockdown cells maintains its helicity, perhaps because the paraflagellar rod remains attached to the cilium (Fig 2), recent studies show they exhibit a significant reduction in cell body rotation [56], suggesting a reduced coupling between cell rotation and ciliary motility in FLAM3 knockdown cells [56, 72].

The effects of increased stiffness and cell rotation that enhance trypanosome swimming motility get neutralized when trypanosome cells have a detached cilium. In such cases, the effects of biological perturbations impair the ciliary beat, and the influence of external perturbations like increased viscosity can become dominant [73]. Here, FLAM3 knockdown causes cells to lose a crosslinking flagellar attachment zone protein that keeps most of the ciliary length section attached to the cell body (Fig 2) [32]. Together, this evidence suggests that the biophysical consequences of dissociating the cilium-cell body complex amplify the effects of various external or biological noise, including waveform modulations caused by dynein light chain knockdowns like ΔLC2. In combination with more straightforward biophysical characterizations of ciliary motility properties, e.g., imaging the shape of the beating waveform is easier when it is not bound to the cell body, these considerations make FLAM3 knockdown trypanosome cells an ideal platform to study regulators of ciliary motility.

### Overexpression of recombinant LC2 is sufficient to cause growth and motility defects

Overexpression of the recombinant TbLC2 in both the wild-type (WT/LC2 OE) and ΔFLAM3/ΔLC2 (ΔFLAM3/Rescued) cells resulted in its localization to the cilium (Fig 2D and S3 Fig), in addition to the cytoplasm (Fig 2A). However, overexpression of the recombinant TbLC2 caused growth and motility defects, including that most WT/LC2 OE cells showed extended ciliary beat reversal (S5 Movie) and only partially restored the growth and motility phenotypes in ΔFLAM3/Rescued cells (Figs 5A, 5B, 7, and 8E). Several factors could contribute to these results, including the effect of the overexpression itself on the overall cell physiology and the effect of the C-terminal chimeric tags, (eGFP, His6, and BCCP), on the secondary structure, electrostatic surface potential distribution, and potential introduction of steric interactions with other axonemal proteins. Overexpression of various other cilium-associated proteins, including the intraflagellar transport kinesin, TbKin2b, and inner arm dynein, IAD-1α, caused abnormalities in the formation of the cilium, the adherence of the cilium to the cell body, and the overall morphology of the cell, all possibly leading to slow growth rate [74]. Moreover, an HA tag on the dynein microtubule-binding domain-associated trypanosome outer arm dynein light chain 1 (LC1) hindered the function of dynein [6]. However, we do not expect such a severe effect of tags on the function of TbLC2 in the outer arm dynein complex because, unlike LC1, it likely does not act directly at the interface between dynein and the microtubule [20], but further analysis on the structural and functional implications of tagging TbLC2 would be needed to rule out that possibility. In any of the possibilities mentioned above, the overexpression of recombinant TbLC2 in ΔFLAM3/ΔLC2 double knockdown cells was still sufficient to at least partially rescue nearly all the phenotypes discussed above.

### *T. brucei* offers an attractive model system for studying mechanisms of ciliary motility in uniciliates

*T. brucei* has emerged as an excellent model organism for studying the cell biophysical aspects of eukaryotic cilia and flagella and molecular biophysics of axonemal dynein-based ciliary motility. Ciliary motility is an intricate part of the parasite’s life cycle, so it provides a platform to understand how a myriad of molecules, including axonemal dyneins and dynein-associated protein regulators, underlie processes like cell division, swimming motility, and ultimately virulence. Trypanosome’s tip-to-base beating mechanism, as well as its genetically accessible nature, make it a particularly compelling model to answer questions fundamental questions about ciliary beating. Moreover, its native attachment to the cell body, and the ability to detach it with FLAM3 (or another flagellar attachment zone protein) knockdown enable one to probe various biophysical aspects of ciliary motility and have those effects amplified (as discussed above). In this study, we provided a quantitative analysis of various facets of cell motility by taking advantage of the differences in the tip-to-base and base-to-tip bending waveform propagation beating modes and FLAM3 knockdowns, which facilitated efficient ciliary waveform tracking and beat mode switching analysis, for example.

## Conclusion

In summary, we identified TbLC2 as a ciliary dynein light chain of the Tctex1/Tctex2 family that is necessary for various aspects of ciliary motility in trypanosome cells. Our work demonstrates that dynein light chain 2 regulation of axonemal dynein causes the modulation of the ciliary beat and ultimately leads to cell-scale trypanosome-specific growth and motility behaviors. Specifically, our results suggest that TbLC2 downregulates and coordinates the underlying activity of axonemal dyneins in the cilium. These molecular mechanisms lead to the stabilization and symmetry of the oscillatory beating waveforms, and the modulation of switching between the reverse and forward ciliary beat modes. Ultimately, TbLC2 helps to maintain the persistent directional motility of trypanosome cells.

The work provides the first evidence that TbLC2 is a key regulator dynein-driven ciliary motility in the disease-causing trypanosome. Further in vivo cell-scale research could elucidate TbLC2’s role in providing trypanosome cells the ability to navigate various microenvironments in the tsetse fly vector [75], avoid the immune system in the mammalian host [5], and cross the blood-brain barrier [2], ultimately causing the most critical stage of disease. Additional in vitro single-molecule research could provide details on the fundamental molecular and biophysical mechanisms of TbLC2-related dynein regulation and flagellar waveform generation. We anticipate that a better understanding of the unique ciliary motility mechanisms in trypanosomes could eventually lead to the development of pan Kinetoplastea therapeutics that target unique aspects of their motility mechanisms.

## Materials and methods

### Sequence-based structural modeling of TbLC2

We identified putative *Trypanosoma brucei* homologs of *Chlamydomonas reinhardtii* LC2 (CrLC2, Cre12.g527750[23]) using protein BLAST (http://www.ncbi.nlm.nih.gov/BLAST/ [24]). We considered proteins with > 70% coverage, Expect values (E value) < 1×10^-5^, and a total alignment score > 50 to be good matches. We confirmed the ciliary localization of identified the proteins in the TrypTag database (tryptag.org) [25, 29]. We performed a multiple sequence alignment of the putative TbLC2 with the *C. reinhardtii* LC2 and its homologs in various other species (S1 Fig) using the multiple sequence alignment tool in Clustal Omega [76]. We performed the color-coding of the sequences based on the level of sequence conservation on Jalview 2.11.1.4 [77] by keeping the conservation index threshold at six [78].

We generated a sequence-based structure homology model of TbLC2 using CrLC2 (PDB ID: 7kzn [20]) as a template in the SWISS-MODEL (https://swissmodel.expasy.org) [30]. The cryo-EM CrLC2 structure lacked 22 N-terminal amino acids, likely due to the flexibility of this domain [20]. Therefore, the results from SWISS-MODEL had a similar N-terminal truncation (Fig 1). We also generated a sequence-based, template-free, AI-derived structural model using AlphaFold [31]. To directly compare with the SWISS-MODEL and cryo-EM structures, we used the same 22 amino acid N-terminal truncation in the sequence for AlphaFold.

We performed the 3D visualization of the modeled structures, the structural alignment, and the root mean square deviation (RMSD) calculations using PyMol (PyMol 2.3.4, Schrodinger, LLC). For optimal alignment of the structures, we used an RMSD outlier rejection cutoff of 2 Å and reported the final RMSD conferring the optimal alignment in the results. We used the Adaptive Poisson-Boltzmann Solver (APBS) [79] in PyMol to calculate the electrostatic potential and map it to the molecular surface. We compared the 22 amnio acid N-terminal truncated AlphaFold structure with the full-length AlphaFold predicted structure (S10 Fig), and we found the differences to be small (root mean square deviation, RMSD = 0.25 Å, S10 Fig).

We determined the residues at the CrLC2-LC9 binding interface by calculating the difference in accessible surface area between the CrLC2/LC9 complex and isolated CrLC2 and CrLC9 proteins (dASA) using PyMol. We set the cutoff for identifying interface residues at 1 Å^2^ (a cutoff of 0 would mean all the residues are included as interface residues).

### Cloning GFP-tagged TbLC2 constructs

We adopted a well-developed strategy to introduce epitope tag-encoding DNA into specific target loci in the trypanosome genome using homologous recombination [74, 80]. We extracted the *T. brucei* 29- 13 genomic DNA (E.Z.N.A Tissue DNA Kit, Omega Bio-Tek, GA) and PCR amplified the full-length coding sequence of the putative TbLC2 gene (Tb927.9.12820, see Results) from the genomic DNA (Table A in S1 Text for primers). We also PCR amplified His6/eGFP (for nickel column affinity and fluorescent localization) and the biotin-binding domain of Chlamydomonas’s biotin carboxyl carrier protein (BCCP) [81, 82] coding sequences (Table A in S1 Text for primers) from the pOCC98 plasmid [83] (gift from Aliona Bogdanova, Max Planck Institute of Molecular Cell Biology and Genetics) and the wt-lc2-bccp plasmid [81] (gift from Ritsu Kamiya, Kyoto University, Kyoto Japan), respectively. We designed all three sets of primers with the NEBuilder Assembly Tool (https://nebuilder.neb.com/) and assembled the PCR products using the HiFi

DNA assembly one-step multiple fragment cloning technique (NEBuilder HiFi DNA assembly cloning kit, E5520, New England Biolabs, MA). We ligated the assembled products into a pXS2 plasmid [84] using the pXS2.Pex13.2 [85] plasmid (gift from Meredith Morris, Clemson University, SC) from which we excised the Pex13.2 fragment with ClaI (R0197S, New England Biolabs, MA) and EcoRI restriction enzymes (R3101S, New England Biolabs, MA. We used the newly constructed pXS2.TbLC2.BCCP.His6/eGFP plasmid as a template to PCR amplify the TbLC2.BCCP.His6/eGFP sequence using TbLC2.BCCP.His6/eGFP Forward and TbLC2.BCCP.HIS6/eGFP Reverse primer set (Table A in S1 Text for primers).

We used pLEW100V5 [80, 86] (a gift from George Cross, Rockefeller University; Addgene plasmid # 24011; http://n2t.net/addgene:24011; RRID:Addgene_24011), a pLEW100 vector derivative, to create an inducible expression vector system with phleomycin resistance, which enabled selection of stable transfectants. The pLEW100V5 plasmid has T7 terminators that regulate the rRNA promoter and drives the extremely high gene expression level. Overexpression of the recombinant TbLC2 facilitated the stable incorporation of the TbLC2.BCCP.His6/eGFP (called TbLC2::eGFP in the Results and Discussion sections) into the axoneme, enabling it to out-compete any endogenous TbLC2 proteins. We excised the luciferase gene and linearized the pLEW100V5 plasmid with HindIII (R3104S, New England Biolabs, MA) and BamHI (R3136S, New England Biolabs, MA) restriction enzymes. We then assembled the PCR amplified TbLC2.BCCP.His6/eGFP dsDNA segment into the linearized pLEW100V5 vector using HiFi DNA assembly to generate the pLEW.TbLC2.BCCP.His6/eGFP plasmid (S11 Fig). The assembly of coding sequences of TbLC2, BCCP, eGFP, and His6 into a single continuous open reading frame ensured the expression of a ∼50 kDa protein structure once induced in trypanosome cells. We verified the correct insertion of LC2::BCCP His6/eGFP into the linearized pLEW100V5 plasmid by agarose gel electrophoresis and Sanger sequencing using the pLEW sequencing primers set (Table A in S1 Text for primers) [87].

While designing the vectors and cloning strategy, we ensured the vectors had only one unique NotI restriction site later used for linearization and stable integration into the rRNA locus [88].

### Cloning RNAi constructs

To knock down the expression of targeted genes in trypanosomes, we used the tetracycline-inducible pZJM RNAi vector [89]. The pZJM vector contains a phleomycin selection marker and a sequence to facilitate the homologous integration of the linearized vector at the silent rRNA spacer locus [89].

To generate the FLAM3 RNAi knockdown plasmid pZJM.FLAM3 (S11 Fig), we excised the tubulin gene from a pZJM.tubulin RNAi plasmid (a gift from James Morris, Clemson University) using XhoI (R0146S, New England Biolabs, MA) and HindIII (R3104S, New England Biolabs, MA) restriction enzymes and inserted a 383 bp fragment of FLAM3 (Tb927.8.4780 [27]) that we PCR amplified (Table A in S1 Text for primers) from the *T. brucei* genomic DNA into the linearized RNAi vector.

To generate the TbLC2 RNAi knockdown plasmid, pZJM.TbLC2 (S11 Fig), we excised the FLAM3 fragment out of pZJM.FLAM3 using XhoI and HindIII restriction enzymes and ligated in a 125 bp fragment of 3’-UTR region of TbLC2 that we PCR amplified from extracted genomic DNA using TbLC2 3’-UTR XH primer set (Table A in S1 Text for primers). We targeted the 3’-UTR region of TbLC2 to ensure that the RNAi only knocks down the endogenous gene and not the expression of recombinant TbLC2::eGFP in the rescue cell lines as the recombinant TbLC2 does not contain its UTRs.

To generate the FLAM3/LC2 double RNAi knockdown plasmid, pZJM.FLAM3.TbLC2 (S11 Fig), we linearized the pZJM.FLAM3 plasmid with the XhoI restriction enzyme and ligated in the LC2 3’-UTR target sequence that we PCR amplified from genomic DNA using TbLC2 3’-UTR XX primer set (Table A in S1 Text for primers). We used calf intestinal alkaline phosphatase (M0290, New England Biolabs, MA) to dephosphorylate the 5’ and 3’ ends of the linearized plasmid to prevent self-ligation of linearized plasmid [90]. By including the RNAi target for both FLAM3 and LC2 in the same plasmid, under the same promotor, we created double knockdown cell lines without performing a double transfection and risking losing one or the other plasmid or requiring another selection marker.

### Maintaining and culturing *T. brucei* cells

We used the procyclic form of *Trypanosoma brucei brucei* (strain 29-13, a gift from James Morris, Clemson University, hereafter called wild-type) for all the genetic modification and epitope tagging. *T. brucei* 29-13 was derived from the 427 strain [91], and it encodes a T7 polymerase (selected for using G418, geneticin, at a final concentration of 15 µg/mL) and a tetracycline repressor (selected for using hygromycin at a final concentration of 50 µg/mL) for induced gene expression [91]. We grew the cells on growth media consisting of SDM-79 [92] (custom produced by Life Sciences Research Products, Thermo Fisher Scientific for the Eukaryotic Pathogens Innovation Center labs, Clemson University, Clemson, SC) supplemented with 10% heat-inactivated tetracycline-free fetal bovine serum (S162TA, Biowest, MO) and porcine hemin at 7.5 µg/mL final concentration (AAA11165-03, Alfa Aesar, MA) to have a population doubling time of 8-9 hours with a logarithmic growth phase between 1×10^6^ and 1×10^7^ cells/mL [93]. We maintained the cells in a humidified environment supplied with 5% CO_2_ and at 27 °C by diluting the culture 10-100-fold every 2-4 days into 5 mL of growth media in 25 cm^2^, 50 mL vented cap culture flasks. We induced knockdown and overexpression in cells transfected with doxycycline-inducible plasmids pZJM and pLEW100V5 plasmid constructs by dosing the cells with 1 µg/mL final concentration of doxycycline.

To assess the effect of physical shaking on cell growth, clustering, and clumping, we split the doxycycline-induced cell cultures into two flasks. We placed one on an orbital shaker (Advanced Dura- shaker, 10159-960, VWR International, LLC) set at 85 rpm within the incubator and cultured the other one under normal conditions, without shaking.

### Transfecting *T. brucei* cells

We transfected plasmids into *T. brucei* cells using electroporation [94] to generate the cell lines used in this study (Table B in S1 Test for cell lines). We maintained the cells at mid log phase (5-10×10^6^ cells/mL) for at least two generations to optimize transfection efficiency. We used a total of 5×10^7^ cells for each transfection and always mirrored the transfection with negative control (with just water instead of DNA) to monitor the action of selection drugs. In short, we linearized 40 µg of the vector with NotI (R0189S, New England Biolabs, MA), mixed the linearized plasmid with 5×10^7^ cells that we had washed with 10 mL of cytomix (120 mM KCl, 25 mM HEPES, 0.15 mM CaCl_2_, 10 mM K_2_HPO_4_, 2 mM EDTA, 5 mM MgCl_2_, titrated to pH 7.6 with KOH) [95] and resuspended to 400 µL into a 4 mm cuvette. We electroporated the cells with two pulses (exponential decay pulse mode, 1500 V, 24 µF) separated by 10 s (Gene Pulser Xcell Electroporation System, Bio-Rad Laboratories, Inc., CA). We added 12 mL of growth media supplemented with 15 µg/mL G418 and 50 µg/mL hygromycin and put 0.5 mL of the electroporated cells per well into a 24-well plate. After incubation for 24 hours, we added 0.5 mL of growth media supplemented with selection drugs (20 µg/mL of blasticidin for pLEW100V5 and 5 µg/mL of phleomycin for pZJM vectors, final concentration) to each well and allowed the cells to grow for 10-14 days at 27°C and 5% CO_2_ and checking the wells every other day for cell growth.

We used a limiting dilution method [96] to generate a clonal cell population from the pool of stable cells by seeding 0.5 cells per well into a 24-well plate containing 400 µL of conditioned growth media (prepared by growing wild-type cells to a density of ∼10^7^ cells/mL in growth media to mimic the environment of a well-grown colony for the growth of single cell in a well) supplemented with the corresponding selection drugs. We induced the clonal cell populations from each well and screened for eGFP expression using flow cytometry, detached cilium using bright field microscopy for RNAi knockdowns, and quantitative reverse transcription PCR (RT-qPCR) for FLAM3 and LC2 knockdowns.

### Flow cytometry

We assessed the fraction of the fluorescent cells in the total cell population, as well as their eGFP expression level, using flow cytometry (CytoFLEX LX, Beckman Coulter, CA, USA). We harvested 1 mL of mid-log phase (5-10×10^6^ cells/mL) wild-type and transgene-induced cells by centrifuging them at 1000×g for 10 minutes. We washed the cell pellets with 1 mL of PBS and resuspended them in 2 mL PBS. We analyzed the GFP fluorescence of ∼20,000 cells using the FITC channel and compared the eGFP fluorescence of cells within an experiment, but not between the experiments conducted at different times.

### Confirmation that RNAi suppresses the expression of TbLC2 using RT-qPCR

The morphological phenotype of the cilium separating from the cell body (Fig 2) simplified our assessment of the FLAM3 RNAi knockdown effectiveness. However, we did not observe morphological differences between ΔFLAM3 and ΔFLAM3/ΔLC2 cells with light (Fig 2C) or scanning electron (Fig 2B) microscopy, leaving open the question of whether the LC2 knockdown was effective. However, our RNAi double knockdown strategy of putting both RNAi targets under the same promotor (S11 Fig) and the extensive detachment of the cilium from the cell body from both the ΔFLAM3 and ΔFLAM3/ΔLC2 cells (Fig 2) suggest that the LC2 RNAi was likely successful, as well.

We confirmed the extent of FLAM3 and endogenous TbLC2 RNAi knockdown using semi-quantitative reverse transcription PCR (RT-qPCR). We isolated the total RNA from 2×10^7^ cells using a Monarch Total RNA Miniprep Kit (T2010, New England Biolabs, MA) as described by the manufacturer with an additional 30-minute, room temperature, in-tube DNAse I (M0303S, New England Biolabs, MA) treatment to eliminate the gDNA contamination. We used the Luna Universal One-step RT-qPCR kit (E3005S, New England Biolabs, MA) to determine the relative abundance of target mRNA between two samples. We prepared 20 µl samples normalized by total RNA content and performed the reverse transcription and cDNA amplification (gene-specific RT-qPCR primer sets, Table A in S1 Text) in a single reaction using a CFX Opus real-time qPCR machine (CFX96, Bio-Rad Laboratories, CA). Briefly, we used a single reverse transcription step (10 min at 55 °C) and an initial denaturation (1 min at 95 °C), followed by 40 cycles of 15 s denaturation and 30 s extension.

We performed semiquantitative RT-qPCR analysis of the threshold cycle, Ct, data to calculate fold change in target mRNA expression, as described previously [97]. We normalized the reactions with total RNA content and compared the cycle threshold between the wild-type and ΔLC2 and ΔFLAM3/ΔLC2 cells. We found a significant reduction in endogenous LC2 gene mRNA in both the ΔLC2 and ΔFLAM3/ΔLC2 cells (reduced to 52.9 ± 2.9% and 43.4 ± 3.8% of the wild-type level, respectively) after 72 hrs of induction (S12 Fig). We found a higher magnitude reduction in the FLAM3 gene mRNA level in ΔFLAM3/ΔLC2 cells (reduced to 16.1 +/- 3.7 % of wild-type) after 72 hrs of induction compared to the ΔLC2 cells (S12 Fig).

### *T. brucei* sedimentation assay

We quantified the motility of *T. brucei* cells by examining their sedimentation rate. Since cells with motility defects tend to settle to the bottom of an undisturbed culture, we quantified the extent of motility defects by measuring the relative decrease in the absorbance at 600 nm (OD600) near the top of a cuvette with cells [98]. We put 500 mL of log phase density (1×10^6^-1×10^7^ cells/mL) cells into 1 mL disposable cuvettes (UVette, 952010051, Eppendorf AG, Hamburg, Germany). We measured the OD600 every hour for 7 hours in each of two cuvettes for each cell line, with one cuvette mixed by pipetting at each measurement and the other kept undisturbed throughout the experiment, using a spectrophotometer (V-1200 Spectrophotometer, 10037-434 VWR International, LLC). To isolate the effect of motility defects from the change in OD600 due to cell growth defects, we calculated the ΔOD600 by subtracting OD600 reading for mixed cuvette from that for stationary cuvette. We performed all sedimentation assays in triplicate.

### Immunofluorescence microscopy

We examined the subcellular localization of eGFP tagged TbLC2 using wide-field fluorescence microscopy. We took all immunofluorescence images using an inverted microscope (Eclipse Ti-E, Nikon Instruments, Inc.) with a Plan Apo 60x water immersion objective. We acquired the images at 16-bit depth and 200-2000 ms exposure time with a CCD camera (CoolSNAP HQ2 Monochrome, Photometrics, AZ). We processed the images and performed quantitative analysis using ImageJ [99, 100].

We cultured eGFP expressing cell lines in induction media to a density of 5×10^6^ cells/mL. We washed the cells with PBS, fixed them on a glass slide using 2% paraformaldehyde, and rinsed them with ample wash solution (0.1% normal goat serum, NGS, in PBS) before applying a 0.1 M glycine solution to quench unreacted aldehydes after fixation and hence reduce background fluorescence caused by aldehydes[101]. After 15 minutes, we rinsed the cells with wash solution and permeabilized them with permeabilization solution (0.5% Triton-x 100 in PBS). We washed the cells with wash solution and added 50 µl of mounting solution (DAPI Fluoromount-G, 0100-20, Southern Biotech, Birmingham, AL). Finally, we sealed the sample with a coverslip and nail polish.

We probed the localization of paraflagellar rod (PFR) in permeabilized, fixed the cells using indirect immunofluorescence tagging [102]. Briefly, we washed the cells with ample wash solution and blocked the sample using blocking solution (10% NGS and 0.1% Triton X-100 in PBS) for 1 hour at room temperature. We incubated the cells with an anti-PFR2 antibody [6] (rabbit anti-PFR2 serum, a gift from Kent Hill, UCLA). After washing the sample with ample wash solution, we incubated the cells with an Alexa Fluor 635 conjugated goat anti-rabbit secondary antibody (A-31577, Thermo Fisher Scientific, Inc., MA) diluted 100-fold in blocking solution for an hour. Subsequently, we washed and mounted cells with DAPI-containing mounting solution and finally sealed them with a coverslip as described above.

### Extraction of cilia

We harvested 1×10^8^ FLAM3 RNAi-induced cells by centrifuging at 1500×g for 10 min at 4°C, washed the pellets in PEME buffer (EGTA 2 mM, MgSO_4_ 1 mM, EDTA 0.1 mM, PIPES free acid 0.1 mM, pH 6.9), and resuspend them into a 50 mL conical tube with 2 mL of PEME. We vortexed the cells for 15 minutes at 3200 rpm and centrifuged them at 420×g in a swinging bucket rotor for 10 minutes. We separated the dissociated cilia from the cell bodies by transferring the supernatant to 1.5 mL microcentrifuge tubes and centrifuging them at 420×g for 5 minutes. We saved the supernatant (cilia extract) and resuspended all the pellets in a 50 mL conical tube with 2 mL of PEME buffer. We performed a second separation by repeating the centrifugation steps. We combined the supernatant-containing ciliary extracts and centrifuged them at 25,000×g for 20 minutes. We stored the cell body and ciliary pellets at -80°C.

We modified a previously published method of cell fractionation to extract the cilia [45] from FLAM3 uninduced or wild-type cells with cilia still intact to the cell body. Briefly, we first harvested and washed 5×10^7^ cells with PEME buffer as described above. We then detergent extracted the cells by resuspending the cell pellet in PEME buffer with 1% IGEPAL (CA-630, Alfa Aesar, MA) and incubating on ice for 15 minutes. We centrifuged the cells at 3400×g for 6 minutes at 4°C and saved the membrane and cytoplasmic content-containing supernatant (S1) and the cilia and subpellicular microtubule corset- containing pellet (P1). We resuspended P1 in PEME buffer with 1% IGEPAL and 1 M NaCl and incubated the mixture for 45 minutes on ice to depolymerize the corset microtubules. We then centrifuged the mixture at 16000×g for 15 minutes at 4°C to separate it into a depolymerized corset microtubule and associated protein-containing supernatant (S2) and an axoneme, paraflagellar rod, and basal body-containing pellet (P2).

### Western blotting

We probed the expression of eGFP tagged TbLC2 in various cell and ciliary fractions using anti-eGFP antibodies and standard immunoblotting methods. Briefly, we generated cell lysates from various cellular and ciliary fractions, incubated them in SDS-sample buffer at 95°C, loaded them into polyacrylamide gel wells and ran them for approximately 45 minutes at 190 V in a mini-PROTEAN tetra vertical electrophoresis cell (Bio-Rad Laboratories, Inc). We transferred the protein bands onto PVDF membranes (170-4156, Bio-Rad Laboratories, Inc) with a Trans-Bolt Turbo System (Bio-Rad Laboratories, Inc), blocked the membrane in blocking buffer (1% skimmed milk powder prepared in PBS with 0.1% Tween), incubated it in mouse anti-GFP primary antibody (SC-9996, Santa Cruz Biotechnology, Inc.) diluted to 1:800 in blocking buffer, washed the membrane thoroughly with PBST (0.1% Tween in PBS), and incubated it in alkaline phosphatase-conjugated goat anti-mouse secondary antibody (31328, ThermoFisher Scientific) diluted to 1:5000 in blocking buffer. We detected the bound alkaline phosphatase labeled antibodies by incubating the membrane in BCIP/NBT (5-bromo, 4-chloro, 3- indolylphosphate/nitro-blue tetrazolium, AMRESCO, LLC.) substrate for 30–45 minutes. Finally, we scanned the membrane (Epson Perfection V700 photo scanner) and quantified the bands with ImageJ [99, 100]. We also probed the samples using rabbit anti-detyrosinated tubulin primary antibody (AB3201, Sigma-Aldrich, MO) diluted to 1:5000 in blocking buffer and alkaline phosphatase-conjugated goat anti-rabbit (31342, Thermo Scientific, MA) secondary antibody diluted 1:7500 in blocking buffer as a loading control.

### Nuclei and kinetoplast characterization

We quantified the number and separation of nuclei and kinetoplasts in cell various cell populations to characterize the defects in the replication and translocation of nuclear and kinetoplast DNA during the cell division. We fixed the cells, stained their DNA using DAPI containing mounting solution (DAPI Fluoromount-G, 0100-20, Southern Biotech, Birmingham, AL), and imaged them using a wide-field fluorescence microscope, as described above. We classified the cells as xKyN, where x is the number of kinetoplasts and y is the number of nuclei present, and x and y = N indicates cells with more than two nuclei and kinetoplasts as previously described [35, 103].

### Electron microscopy

We imaged the structure of the cilia with negatively stained electron microscopy. We fixed the cells by incubating 5 mL of culture with electron microscopy grade glutaraldehyde (16300, Electron Microscopy Sciences, Inc., PA) at a final concentration of 2.5% at room temperature for 10 minutes followed by and an overnight fixation with Karnovsky’s fixative reagent (2.5% glutaraldehyde, 2% paraformaldehyde, and 0.1% tannic acid prepared in 0.1 M phosphate buffer, pH 7, 15720, Electron Microscopy Sciences, Inc., PA). We washed the cells once with 0.1 M pH 7.2 phosphate buffer, resuspended them in 0.1 M pH 7.2 phosphate buffer with 0.02% sodium azide, and stained them with 1% osmium tetroxide before running the sample through a graded ethanol series to dehydrate the samples. For TEM, we infiltrated the ethanol-soaked samples with LR white embedding resin and cured them for 24 hrs in a 60°C oven. We sectioned the samples with a microtome, placed thin sections on copper grids, and captured the transmission electron brightfield micrographs with a Hitachi SU9000 UHR operated at 100 kV. For SEM, we soaked the dehydrated sample in 1:1 ethanol:HMDS (hexamethyldisilazane, 16700, Electron Microscopy Sciences, Inc., PA), followed by pure HMDS. We air-dried the samples overnight, placed them onto silicon wafers, and sputter coated them with platinum. Once dry, we imaged the samples using SEM (Hitachi SU5000 at an accelerating voltage of 2kV).

### Motility traces

We tracked the motility of trypanosome cells using widefield light microscopy and quantitative image analysis. We prepared a 100 – 150 μm deep motility chamber constructed from parafilm (PM-999, Bemis, WI) sandwiched between easy cleaned [104] glass slides and No. 1.5 22 mm x 22 mm cover glasses (16004-302, VWR). We passivated the chamber by incubating it with 0.25% poly-L-glutamate (P4886, Sigma-Aldrich, Inc.) [57] prepared in PBS for 25 minutes and washing out the excess poly-L-glutamate with distilled-deionized water and ethanol. We diluted cultured and induced cells to a final density of 1×10^6^ cells/mL using fresh growth media and equilibrated them to room temperature. We transferred 10 µl into the motility chamber, sealed it with the nail polish, and imaged it with phase-contrast microscopy (Aus Jena Telaval 3 inverted binocular microscope) at 5x magnification (Aus Jena planachromat, NA 0.1). We recorded movies at 45 frames per second using a CCD camera (Grasshopper3 USB3 GS3-U3-15S5M-C, Teledyne FLIR, LLC.) for 10 seconds.

We preprocessed the image sequences in ImageJ [99,100,105] by subtracting the maximum projection from each frame. We thresholded the images and applied a spot-enhancing 2D filter (SpotTracker plugin [106, 107]) to further enhance the cell spot and reduce the background noise. We cropped the frames to include only single cells. We quantified motility by tracking single, non-intersecting cell paths to minimize the breaking of tracks using the wrMTrck [108] and ParticleTracker [109] plugins with the following parameters: a) minSize of 100 pixels^2^, b) maxSize of 400 pixels^2^, and c) threshMode Otsu for wrMTrck and a radius of 15 pixels and cutoff of 3 pixels for ParticleTracker. We tracked cells over the entire sequence of frames (10 s) to minimize bias due to inequality in tracked time intervals [54].

We used DiPer [54] to calculate the directional ratio, curvilinear speed, and velocity of the tracked trypanosomes. Briefly, directional ratio (DR) as a function of time is the magnitude of the total straight- line vector displacement divided by scalar distance (integrated path length) covered by cell from the initial time point at each time point. We used the DR at the last time point of each track to classify the cell population into three groups: A. high persistence (DR > 0.2, most persistent swimmers), B. medium persistence (0.05 < DR < 0.2, persistent swimmers with intermittent tumbling), and C. low persistence (DR < 0.05, tumblers) (S2 Movies for examples of each). Curvilinear speed is the time average of an individual cell’s instantaneous speed. Velocity is the magnitude of the total straight-line vector displacement between the two ends of the tracks divided by the 10 s time length of the track.

### Optical tweezer assay and power spectral density analysis

We used a custom-built, single-beam optical tweezer to trap the cells and measure the fluctuation of the transmitted laser intensity in the back focal plane of the condenser due to the scattering of the laser caused by cell displacements driven by rotation and ciliary beating. Optical tweezer-based frequency analysis has multiple advantages over the quantitative image analyses of high-speed video microscopy alternatives, including fewer constraints on cell rotations (S1 Movies) and out-of-plane beating waveforms than occur in thin (<10 μm) imaging chambers used for microscopy, the ability to probe higher order frequencies and get better resolution with power spectral density analysis of data collected at 10 s of kHz, and higher throughput than having to perform time-consuming video analyses. Briefly, the optical tweezers use a 1064 nm, 10 W ytterbium fiber laser (YLR-10-1064-LP, IPG Photonics) to generate the trapping laser beam focused into the sample plane using a CFI Plan Apochromat Lambda 60x N.A. 1.4 oil immersion objective lens (Nikon Instruments, Inc.).

We harvested cultured and induced cells in the log growth phase and diluted them to a final concentration of 1×10^6^ cells/mL in fresh growth media. We flowed the cells into a 2-3 mm wide chamber constructed as described above, sealed the chamber with nail polish, and allowed the media to equilibrate to room temperature for 10-15 minutes. We trapped the cells at minimal laser power at the sample plane (10-25 mW) to maintain cell viability by minimizing photo and thermal damage [47]. We captured the time series laser fluctuation data with a quadrant photodiode (QPD, QP45-Q HVSD, First Sensor Inc.) using back focal plane detection [110] at 500 kHz and smoothened by averaging to a final rate of 25 kHz using an FPGA (PXI-7854R, NI) and custom-written LabView (NI) VIs. We calculated the power spectral density (PSD) from the time series data using the FFT Power Spectrum and PSD built-in LabView functions and averaged three such PSDs for each measurement. We used the QPD sum signal fluctuation (represents position fluctuation axially [111]) and difference signal fluctuation (represents position fluctuation in the x-y plane [111]) for determining the ciliary beat and cell rotation frequencies, respectively, in the PSD plot.

### High-speed imaging, shape tracking, and measuring curvature and ciliary beat switch

We recorded high-speed image sequences in an assay chamber constructed using microscopic beads as spacers as previously described [104], with minor modifications. In brief, we 10-fold diluted a 5% w/v stock of polystyrene beads (PP-50-10, Spherotech, Inc.) and put 5 µm onto an easy cleaned [104] microscope glass slide. We placed an 18 mm x 18 mm easy cleaned coverslip on the bead solution droplet and sealed the sides of the coverslip (perpendicular to the length of the slide) with nail polish. We washed the beads out with distilled deionized water using vacuum suction. The chambers had a depth of 5.65 +/- 0.12 µm (as measured by focusing the objective at the two inner surfaces of the chamber, mean +/- SEM, N=16), which was larger than the cell width (1.54 – 3.5 µm [112]) but smaller than the cell length (S5 Fig) and thus confined the cell body to the focal plane while allowing for free ciliary beating.

We harvested cultured and induced cells in the log growth phase and diluted them to a final concentration of 1×10^6^ cells/mL in fresh growth media. We sealed the chamber with nail polish and immediately imaged the cells with bright field illumination using a CFI Plan Apochromat Lambda 60x N.A. 1.4 oil immersion objective lens (Nikon Instruments, Inc.). We captured 3-second image sequences at 200 frames per second with a minimum exposure time (250 µs, to minimize motion blur) using a high-speed CMOS camera (M-PRI-1000, AOS Technologies AG). We slightly defocused the sample stage to facilitate the shape tracking.

As explained in the motility tracking section above, we preprocessed the image sequences in ImageJ [99, 100]. We modified custom-written MATLAB scripts [113] (The MathWorks, Inc.) to track individual beating cilia and calculate the tangent angle with respect to a horizontal reference axis, *ψ*, at equally spaced points along the ciliary arc length, *s* (Veikko Geyer and Benjamin Friedrich provided the source code for both scripts).

To determine the ciliary beat amplitude (A), we first subtracted the average ciliary shape from the tangent angles measured at each point along s and calculated the amplitude as the half-width of the waveform shape formed by the tangent angle distributions [59]. We then measured the ciliary beat wavelength as twice the separation, in terms of arc length normalized to the contour length (s/L), between the maximum and the minimum tangent angles. We calculated the curvature as a function of 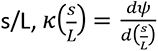. We calculated the curvature at 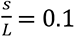, as measured from the cilium’s tip, and used the absolute values of the curvature (to prevent the oppositely directed curvatures canceling each other) to calculate the time average curvature-at-the-tip.

We calculated the maximum curvature asymmetry ratio, *AR*_MC_, by dividing the maximum curvature found in each opposing bend direction as a single ciliary bend wave propagated along the cilium. Since we have no marker for cell or cilium orientation, we calculated the asymmetry ratio such that *AR*_MC_ ≥ 1. An *AR*_MC_ = 1 indicates that the bending wave propagated symmetrically along the cilium, and an *AR*_MC_ > 1 indicates that the bending wave propagated asymmetrically along the cilium.To determine the fraction of time the cells spend in the tip-to-base (regular) and base-to-tip (reverse) ciliary beating modes, we analyzed the ciliary beating of each cell, frame by frame, over the entire period of 3 seconds (600 frames). We noted the reverse beat dwell time as the interval between the initiation of base-to-tip ciliary bend propagation (identified by the traveling of the ciliary bend initiated at the ciliary base towards the tip over the 600 frames) to the initiation of the next regular ciliary bend at the tip and divided the dwell time by 3 seconds (or 600 frames).

## Author contributions

**Conceptualization:** Joshua Alper.

**Data Curation:** Subash Godar, Joshua Alper.

**Formal Analysis:** Subash Godar, Joshua Alper.

**Funding Acquisition:** Joshua Alper.

**Investigation:** Subash Godar, James Oristian, Valerie Hinsch, Katherine Wentworth, Ethan Lopez, Parastoo Amlashi, Gerald Enverso, Joshua Alper.

**Methodology:** Subash Godar, James Oristian, Valerie Hinsch, Katherine Wentworth, Ethan Lopez, Parastoo Amlashi, Gerald Enverso, Samantha Markley, Joshua Alper.

**Project Administration:** Joshua Alper.

**Resources:** Subash Godar, James Oristian, Valerie Hinsch, Katherine Wentworth, Gerald Enverso, Samantha Markley, Joshua Alper.

**Software:** Subash Godar, Joshua Alper.

**Supervision:** Joshua Alper.

**Validation:** Subash Godar, Joshua Alper.

**Visualization:** Subash Godar, Joshua Alper.

**Writing – Original Draft Preparation**: Subash Godar, Joshua Alper.

**Writing – Review & Editing**: Subash Godar, Joshua Alper.

## Acknowledgments

This work was supported by the National Institute of Allergy and Infectious Diseases (NIAID) of the National Institutes of Health under award number R15AI137979 and the National Institute of General Medical Sciences (NIGMS) of the National Institutes of Health under award number P20GM109094. Additional funding was provided by the Clemson University Division of Research, College of Science, Department of Physics and Astronomy, Creative Inquiry program, and University Professional Internship and Co-op Program. We are grateful to the Clemson Light Imaging Facility, Terri Bruce, and Rhonda Reigers Powell for the use of and their help using the light microscopes. We are also grateful to the Clemson Electron Microscopy Facility, Laxmikant Vyankatesh Saraf, and George Wetzel for the use of and their help using the TEM and SEMs. We also acknowledge Aliona Bogdanova (Max Planck Institute for Molecular Cell Biology and Genetics), Jim Morris (Clemson University), members of the J. Morris Lab, Meredith Morris (Clemson University), Kent Hill (UCLA), and Ritsu Kamiya (Kyoto University) for various reagents and samples, use of their equipment, and general advice. We also thank Veikko Geyer and Benjamin Friedrich for the MATLAB source code we adapted to track beating trypanosome cilia. Finally, we are thankful for Marija Zanic, her careful reading of the manuscript, and many fruitful discussions.

## Supporting information captions

**S1 Text. Supporting Tables A and B.**

S1 Fig. Multiple sequence alignment of putative Tctex-type dynein light chains in the *Trypanosoma brucei* genome and LC2 homologs from other genomes. *T. brucei* A and B represent the two closest hits for the *C. reinhardtii* homolog of LC2 (Tb927.9.12820 and Tb927.11.7740, respectively). The color bands indicate the level of sequence conservation, with red indicating identical residues, and the spectrum between orange and yellow indicates level of sequence conservation from high to low. White residues are not conserved. (Materials and Methods).

**S2 Fig. Sequence alignment of TbLC2 (*T. brucei* A) and the *C. reinhardtii* homolog of LC2 (*C. reinhardtii*).** Conservation of hydrophobicity (*red*) and polarity (*blue*) are indicated. The spectrum from light red to dark red indicates lower to higher hydrophobicity, and the spectrum from purple to blue indicates low to high polarity (charge). The *C. reinhardtii* LC2 residues above the black lines are in the LC9 binding interface as determined by using the change in the accessible surface area (dASA) in PyMol (PyMol 2.3.4, Schrodinger, LLC).

**S3 Fig. Multi-histogram plot of eGFP fluorescence intensity measured by using flow cytometry for TbLC2::eGFP overexpressed cell lines.** The frequencies of occurrence are normalized to the mode to facilitate comparisons across cell lines with different cell counts registered by the flow cytometer. The wild-type cell line was used as a control. Cell counts are N= 517500, 61144, and 130600 for WT, ΔFLAM3/Rescued, and WT/LC2 OE cells, respectively.

**S4 Fig. Western blot analysis of LC2::eGFP**. Anti-GPF antibodies stained LC2::eGFP in uninduced (-dox) and induced (+dox) whole-cell (WC), cell body only (CB), and ciliary (cilia) fractions. We used tubulin as a loading control. We extracted the cilia using mechanical shearing and loaded an equal number of cells in all lanes except in the ciliary fractions, which contains cilia extracted from 15-fold as many cells as used in the other lanes. We used tubulin (bottom) as a loading control.

**S5 Fig. FLAM3 knockdown and TbLC2 overexpression cause shorter *T. brucei* cilia.** Length distributions of cilia from strains as indicated. The black lines indicate the mean length, and shaded regions around the line indicate the SE of the mean (N = 76, 76, 52, and 101 for WT, ΔFLAM3, ΔFLAM3/ΔLC2, and ΔFLAM3/Rescued cells, respectively). *** represents *p*-value < 0.0001 and ** represents *p*-value < 0.001.

**S6 Fig. Shaking does not rescue the cell growth defects in LC2 knockdown cells.** Growth curves for ΔFLAM3/ΔLC2 (*purple*) and ΔFLAM3/Rescued (*orange*) cell lines, comparing the effect of mechanical agitation. Shaken (*dashed lines*) and not shaken (*solid lines*) indicate incubation of cell culture with or without orbital shaking, respectively at

90 rpm for the duration of incubation. Points represent the mean of three cultures, and error bars represent the standard error of the mean.

**S7 Fig. Tangent angle characterization of the ciliary beat waveform in FLAM3 knockdown cell derivatives.** Example tangent angle (ψ(s/L)) plotted as a function of arc length (s) normalized to the contour length (L) along the cilium for a typical tip-to-base wave propagation in ΔFLAM3 cells (*left*), and the same tangent angle plotted after subtraction of the average ciliary beating shape (*right*). Data from an example frame (*green*) used to calculate the beat amplitude and wavelength (*black arrows*) and the average shape (the static component of the waveform, *red*) are highlighted both before (*left*) and after (*right*) the average shape subtraction. This cilium exhibits only slightly non-zero mean curvature because the average shape of the ciliary beating waveform has only a slightly non-linear time-averaged tangent angle (*red, left*). This cilium exhibits an approximately symmetric dynamic curvature because the time-averaged tangent angle subtracted waveform (e.g., *green*, *right*) has approximately equal positive and negative maximum curvature. s/L of zero represents the ciliary base in both plots.

**S8 Fig. Curvature during the highly asymmetric reversed base-to-tip beating in FLAM3/LC2 double knockdown cell.** The curvature normalized to contour length (κ/L) plotted as a function of normalized arc length (s/L) along the cilium for a typical highly asymmetric base-to-tip wave propagation in ΔFLAM3/ΔLC2 cells. s/L of zero represents the ciliary base, and the colors (*red* to *magenta*) represent time progression during the propagation of a ciliary bend in a single ciliary beat period. The t = 0 beat profile (*red*) represents ciliary bend initiation towards the base (left side), followed by the successive profiles representing the propagation of the bend towards the tip (*right side*).

**S9 Fig. The static component of ΔFLAM3/ΔLC2 cells ciliary beating waveform**. The mean tangent angle was calculated as a function of normalized arch length (s/L) from the average shape of the beating waveform for forward (tip-to-base) and reverse (base-to-tip) beating waveforms of ΔFLAM3/ΔLC2 cells. Each point represents the mean of the average tangent angle from 8 cells, and the error bars represent the standard error of the mean.

**S10 Fig. Full-length TbLC2 has a high degree of structural conservation with the N-terminal truncated TbLC2 protein, as calculated by AlphaFold**. The template-free model of TbLC2 with the 22 N-terminal amino acids truncated (*left*) to match the resolved residues in the cryo-EM (Fig 1B), and the template-free model of full-length TbLC2 (*right*), both as calculated by AlphaFold.

**S11 Fig. Vector maps of overexpression and RNAi plasmids used.** Vector maps of the tet-inducible pLEW.TbLC2::BCCP His6/eGFP plasmid generated by cloning the preassembled TbLC2, BCCP, His6, and eGFP genes into the ORF of pLEW100V5 plasmid by excising the luciferase gene (*gray*). The expression of the TbLC2::BCCP His6/eGFP gene is driven by the strong rRNA promoter, whereas the T7 promoter drives the expression of the selection marker (blasticidin). The rRNA spacer acts as a target locus during transfection. Vector map of the pZJM.FLAM3 RNAi vector generated by replacing the tubulin sequence (*gray*) between XhoI and HindIII restriction sites with the FLAM3 sequence. Vector map of the pZJM.TbLC2 RNAi vector generated by replacing the FLAM3 (*gray*) sequence between XhoI and HindIII restriction sites with TbLC2 3’-UTR. Vector map of the pZJM.FLAM3.TbLC2 RNAi vector generated by inserting the TbLC2 3’-UTR sequence at the XhoI restriction site. All the RNAi vectors are integrated into the trypanosome genome at the rDNA site.

**S12 Fig. mRNA expression level of FLAM3 and LC2 genes in ΔLC2 and ΔFLAM3/ ΔLC2 cell lines determined by semiquantitative RT-qPCR**. The percentage reported is relative to the expression of the same genes in the WT cells. The error bars represent SEM calculated from triplicates of the experiment performed on the total RNA sample purified.

**S1 Movies. Trapped cells displayed unconstrained-like ciliary beating behavior. A.** Movie of an uninduced cell trapped by the optical tweezer 70-100 µm above the surface of the motility chamber. The cell is free to rotate and maintain the out-of-plane nature of its beating waveform. **B.** Movie of an uninduced cell trapped by the optical tweezer approximately 3 µm above the surface of the motility chamber. The cell is constrained by the glass surface. However, the rotation and out-of-plane beating is observable. The powe spectra used to quantify the beat frequency in Fig 6 were obtaind with cells trapped 70-100 µm above the surface of the surface, like **A.** Both movies recorded using wide-field microscopy with a 60x objective at 45 fps and played back at the same frame rate.

**S2 Movies. LC2 knockdown cells exhibit three distinct categories of directional motility. A.** High, **B.** medium, and **C.** low directional persistence motility. All three movies were recorded using phase-contrast microscopy with a 10x objective at 45 fps and played back at the same frame rate. The scale bar represents 10 μm in each movie.

**S3 Movies. ΔFLAM3/ΔLC2 double knockdown cells show altered ciliary beating leading to a frequent change in swimming direction. A.** A high-speed movie of ΔFLAM3/ΔLC2 double knockdown cells showing a highly asymmetric forward (tip-to-base) beating with an intermittent reversal in ciliary beat direction—both of the phenomena leading to cell reorientation. The movie was recorded at 200 fps with an exposure time of 250 µs and played back at 45 fps. **B.** A typical freely swimming ΔFLAM3/ΔLC2 cell showing frequent reversals in ciliary beat mode and futile swimming motility with low directionality. The movie was recorded at 45 fps and played back at the same frame rate. The scale bar = 10 µm in both movies.

**S4 Movie. ΔFLAM3/ΔLC2 double knockdown cells exhibit ciliary beating locked to a reverse beating over an extensive period showing a longer dwell time in reverse beating mode.** The movie was recorded using a high-speed CMOS camera and a 60x objective at 200 fps with an exposure time of 250 µs and played back at 45 fps. Scale bar = 10 µm.

**S5 Movie. The trypanosome base-to-tip beating mode is not exclusively asymmetric.** Movie of a typical WT/LC2 OE cell exhibiting extensive, a largely symmetric, base-to-tip (reverse) beating mode. This shows the effect of LC2::eGFP overexpression on trypanosome cell motility. The movie was recorded using phase-contrast microscopy at 45 fps and played back at the same frame rate. Scale bar = 10 µm.

**S1 Files. Sequence-based modeled structures (.pdb) of TbLC2. A.** Homology modeled structure of TbLC2 using CrLC2 as a template in SWISS-Model. **B.** Template-free model of TbLC2 built using AI-based AlphaFold modeling with the 22 amino acid N-terminal truncation. **C.** Template-free model of the full-length TbLC2 built using AI-based AlphaFold modeling.

